# *Drosophila* USP22/non-stop regulates the Hippo pathway to polarise the actin cytoskeleton during collective border cell migration

**DOI:** 10.1101/2020.07.02.177170

**Authors:** Hammed Badmos, Neville Cobbe, Amy Campbell, Daimark Bennett

## Abstract

Polarisation of the actin cytoskeleton is vital for the collective migration of cells *in vivo*. During invasive border cell migration in *Drosophila*, actin polarisation is directly controlled by Hippo pathway components, which reside at contacts between border cells in the cluster. Here we identify, in a genetic screen for deubiquitinating enzymes involved in border cell migration, an essential role for non-stop/USP22 in the expression of Hippo pathway components *expanded* and *merlin*; loss of *non-stop* function consequently leads to a redistribution of F-actin and the polarity determinant Crumbs, loss of polarised actin protrusions and premature tumbling of the border cell cluster. Non-stop is a component of the Spt-Ada-Gcn5-acetyltransferase (SAGA) transcriptional coactivator complex, but SAGA’s histone acetyltransferase module, which does not bind to *expanded* or *merlin*, is dispensable for migration. Taken together, our results uncover novel roles for SAGA-independent non-stop/USP22 in Hippo-mediated collective cell migration, which may help guide studies in other systems where USP22 is necessary for cell motility and invasion.

## Introduction

Tightly regulated cell migration is vital for normal development and aberrant migration is involved in a number of human diseases, including tumour invasion and cancer metastasis, inflammatory diseases, and various birth abnormalities (Schumacher, 2019; Stuelten et al., 2018). In many instances, cells move by the process of collective migration *in vivo*, whereby migratory cells remain connected by cell-cell junctions, show group polarisation and coordinated cytoskeletal dynamics (Haeger et al., 2015; Mishra et al., 2019; Norden and Lecaudey, 2019). This mode of migration is exemplified by the movement of border cells in *Drosophila* (video S1). In this process, a cluster of five to eight cells are recruited from the follicular epithelium in the ovary by a pair of non-motile polar cells. Both cell types migrate as a cluster from the anterior basal lamina of the egg chamber, invading the underlying germ line, to the anterior border of the oocyte where they are involved in patterning prior to egg fertilization (Montell et al., 2012).

Studies of this process over the past 20 years have identified key features of the genetic programme required for border cell migration, which control the specification of the migratory cluster (Bai et al., 2000; Montell et al., 1992; Silver and Montell, 2001), organisation of cluster polarity and detachment from the epithelium (Abdelilah-Seyfried et al., 2003; McDonald et al., 2008; Pinheiro and Montell, 2004), timing of migration (Godt and Tepass, 2009; Jang et al., 2009), adhesion of the cluster (Cai et al., 2014; Niewiadomska et al., 1999) and guidance to the oocyte (Bianco et al., 2007; Duchek and Rorth, 2001; Duchek et al., 2001; McDonald et al., 2003). Details have also emerged regarding the dynamic organisation of the actin cytoskeleton which is an essential driver of this process (Plutoni et al., 2019), with recent studies identifying an important role for the Hippo pathway in linking determinants of cell polarity with polarisation of the actin cytoskeleton in migrating clusters (Lucas et al., 2013). Our understanding of the interplay between polarity determinants and the actin cytoskeleton however remain incomplete, as does knowledge of the regulatory networks responsible for first establishing this polarity.

Ubiquitination of proteins by ubiquitin E3 ligases and removal by deubiquitinating enzymes (DUBs) plays important roles in regulating a raft of intracellular functions from protein stability and enzyme activity to receptor internalization and protein-protein interactions (Clague et al., 2013; Swatek and Komander, 2016). There is a growing body of evidence that ubiquitination plays roles in regulating the motility of single cells in culture (Cai et al., 2018), but little is known about its contribution to collective migration *in vivo*. Here we report our identification of *non-stop* (*not*) from a screen of DUBs involved in border cell migration. *not* encodes the USP22 orthologue in *Drosophila* (Martin et al., 1995), and is best known as the enzymatic component of the histone H2B DUB module of the SAGA transcriptional coactivator complex (Koutelou et al., 2010; Lee et al., 2011; Zhang et al., 2008). Histone modifications such as acetylation and ubiquitination are known to modulate the accessibility of genomic loci to transcriptional machinery, with ubiquitination being associated with both activation and repression (Weake and Workman, 2008). Correspondingly, SAGA is associated with the enhancers, promoters and sites of paused RNA polymerase II at genes in multiple tissues during Drosophila embryogenesis, and the Non-stop activity within SAGA is required for full expression of tissue-specific genes (Weake et al., 2011).

Previous work has revealed essential roles for non-stop/ USP22 during embryogenesis in *Drosophila* and mammals (Li et al., 2017; Lin et al., 2012), as well as in neural development (Weake et al., 2008) and lineage specification (Kosinsky et al., 2015). In the *Drosophila* nervous system, loss-of-function mutations in *non-stop* are associated with defects in the migration of a subset of glial cells to their appropriate position in the developing optic lobe and subsequent targeting of photoreceptor axons in the lamina (Martin et al., 1995; Poeck et al., 2001). The underlying mechanisms are not fully understood, but it has recently been suggested that this role may be mediated in part by a SAGA- independent role of Not in deubiquitinating and stabilising the actin regulator Scar (Cloud et al., 2019). Here we find that, in collective border cell migration, *not* functions independently of both Scar and SAGA to regulate the expression of two upstream components of the hippo pathway, resulting in the loss of F-actin polarity, the mislocalisation of polarity determinants, a change in the size and orientation of cellular protrusions and the loss of polarised migration, placing *non-stop* at the top of a regulatory network underlying collective migration.

## Results

### *non-stop* is required for invasive border cell migration

We identified the *Drosophila* USP22 homologue *non-stop* in an RNA interference (RNAi) screen for deubiquitinases (DUBs) required for border cell migration. Wild-type border cell clusters normally reach the oocyte by stage 10 of oogenesis, whereas expression of transgenic inverted repeat constructs for *non-stop* (*not^IR^*) in the outer border cells severely delayed border cell migration (mean percentage migration of the distance to the oocyte ±SEM was 2.5 ±2.5%, n=40, Student’s t-test, *P*<0.0001) (Fig.1A-E). These migration defects could be significantly rescued by a full-length synthetic RNAi-resistant transgene (*not^+r^*, see Methods) confirming the requirement for *non-stop* in migration (Fig.1D,E). Incomplete rescue is most likely an indication of some off-target effects of *not^IR^*. Expression of *not^+r^* alone in both polar and outer border cells had no effect on migration (Fig.1C,E). To further confirm the requirement for *non-stop* in border cell migration, we generated homozygous clones for an amorphic *non-stop* mutant allele (*not^1^*). Notably, border cell clusters genetically mosaic for *not^1^* showed greatly retarded migration, with the severity of the effect being dependent on the proportion of mutant cells in the cluster (Fig.1F-H). Mean migration was reduced by 61.3 ±2.9% (*P*<0.0001, n=101) in clusters containing >50% *non-stop* mutant cells compared to clusters with control clones, where cells migrated normally; these defects were almost fully rescued by transgenic expression of *not^+r^* (Fig.1H). Unlike in controls, splitting of border cell clusters was also observed in 28% of stage 9 or 10 *non-stop* mutant egg chambers (n=138) (Fig.1I-J), indicative of a defect in maintaining the integrity of border cell-border cell contact. Taken together, these data identify *non-stop* as a novel regulator of border cell migration.

**Fig. 1.**
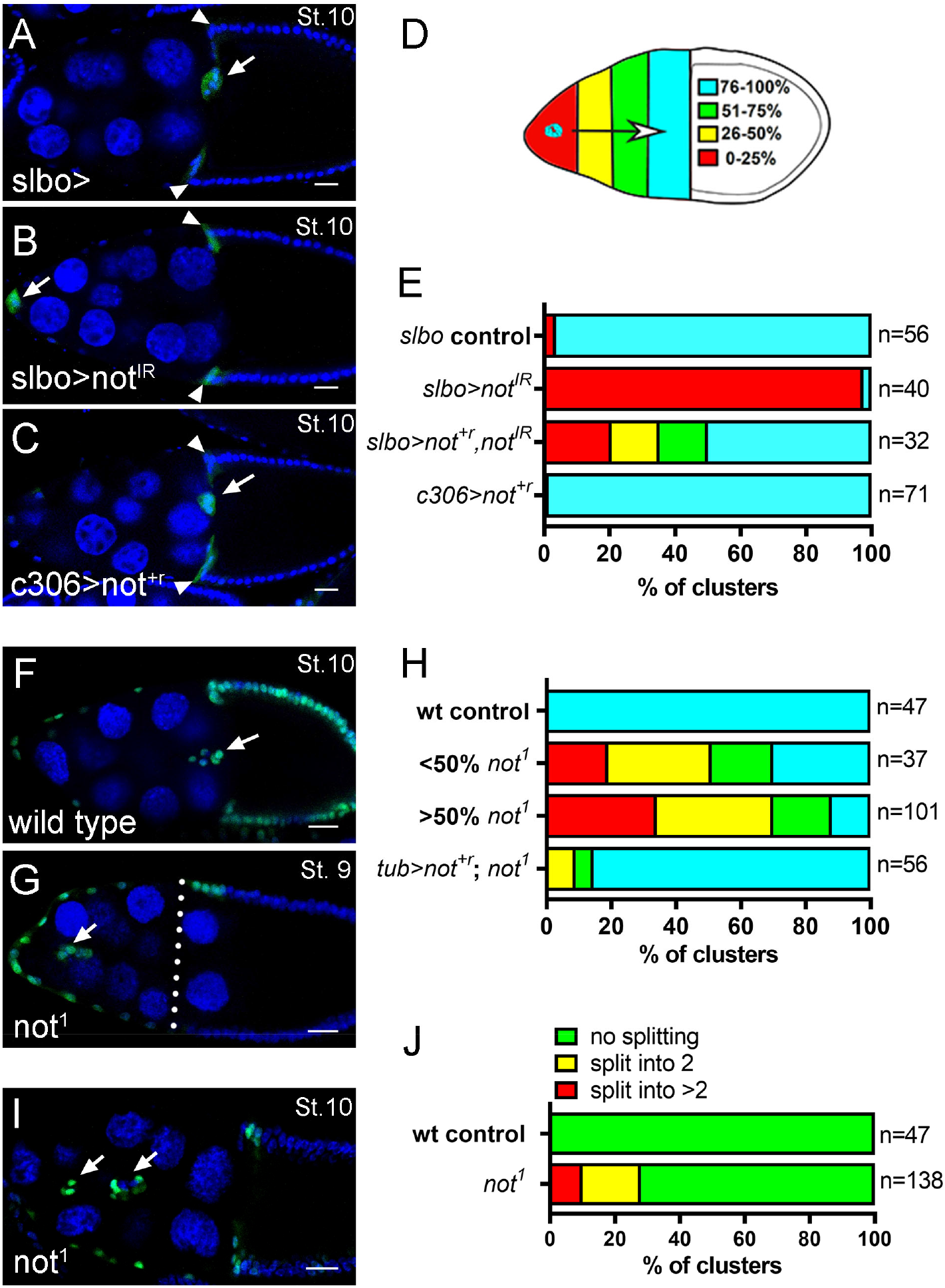
*non-stop* is required for invasive border cell migration. **A-D**, Confocal micrographs of egg chambers at stage 10 labelled with GFP (green) under the control of *slbo-GAL4* to mark border cells (arrows) and TOPRO-3 (blue) to stain all nuclei. Anterior is left, posterior is right. Some GFP expression is also evident in centripetal follicle cells (arrowhead). Bars, 25μm. **A**, Image of *slbo-GAL4* control (slbo>) showing complete migration of the border cell cluster. **B**, RNAi knockdown of *not* under the control of *slbo-GAL4* (slbo>not^IR^) abrogates border cell migration. **C**, In contrast, overexpression of *not^+r^* in the whole border cell cluster using *c306-GAL4* (c306>not^+r^) did not affect migration, indicating *non-stop* is not limiting for migration. Clusters expressing *not^+r^* with *Slbo-GAL4* also migrated normally (not shown). **D**, Migration index for quantification of border cell migration at stage 10, see Methods. **E**, Stacked bar chart summarising migration defects in the indicated genotypes (n= number of egg chambers). The effect of *not* RNAi knockdown can be partially rescued by transgenic overexpression of RNAi-resistant *not* (slbo>not^+r^ *not^IR^*). **F-G,** Confocal micrographs of egg chambers labelled with GFP (green) to mark clones of cells induced with the MARCM technique and TOPRO-3 (blue) to stain all nuclei. Bars, 25μm. Compared to control clones, which routinely complete migration at stage 10 (**F**), *not^1^* mutant border cell clusters display defective migration, with clusters lagging behind overlying centripetal cells (position marked with dotted line) at stage 9 (**G**). **H**, Quantitation of migration defects at stage 10, reveal that the clusters containing >50% mutant cells are more severity affected than those with <50% mutant cells in the cluster; migration is largely restored by *not^+r^* overexpression (tub>not^+r^; not^1^). n= number of egg chambers. **I,** Stage 10 egg chamber showing splitting of *not^1^* mutant border cell clusters; 18% of clusters displayed splitting into two groups of cells, 10% of clusters split into >2 groups of cells (**J**, frequency of cluster splitting, n= number of egg chambers).

### *non-stop* regulates polar cell number

At stage 8 of oogenesis, a pair of anterior polar cells secrete Unpaired (Upd) ligand, which activates the JAK-STAT (Janus kinase-signal transducer and activation of transcription) signalling pathway in surrounding follicle cells, leading to the recruitment of 5-8 follicle cells into a migratory cluster (Beccari et al., 2002; Silver and Montell, 2001). To explore the requirement for *non-stop* in border cell signalling we looked at the expression of *slbo*, a downstream target of Upd-JAK/STAT signalling in the migratory outer border cells, which induces the expression of genes required for migration. The level of a transcriptional reporter, *slbo-lacZ*, was not significantly different between *not^1^* mutant cells and their wild type siblings within mosaic border cell clusters (Fig.2A-C; arbitrary units, mean intensity ±SEM was 47 ±5.3 for *not^1^* n=21, compared to 56 ±5.5 for controls n=23, *P*=0.22). *non-stop* was also not required for the expression pattern of Eyes absent (Fig.2D,E), which is expressed in outer border cells to repress polar cell fate in these cells (Bai and Montell, 2002). Therefore, we conclude that *non-stop* does not affect the expression levels of genes in migratory outer border cells that specify their fate. When we looked at upstream signalling using a *upd-lacZ* reporter (Fig.2F-H), we observed that 32% of *not^1^* mutant clusters possessed more than two *upd-lacZ* positive polar cells (Fig.2I; mean ±SEM was 2.47 ±0.12, n=53), suggesting that some *not^1^* polar cells continue proliferating after stage 2 of egg chamber development when divisions would normally cease (Margolis and Spradling, 1995). *not^1^* clusters also contained on average a 1.7-fold higher number of border cells than controls (Fig.2I; mean ±SEM was 11.1 ±0.2 for *not^1^* n=138, compared to 6.4 ±0.12 for controls n=47, *P*<0.0001) and this was correlated with the number of *upd-lacZ* positive polar cells (Fig.2J; multiple regression R^2^=0.54, n=53, *P*<0.0001), suggesting the presence of additional polar cells led to the recruitment of additional border cells into the cluster. Clusters with more than two polar cells had a significantly reduced degree of migration compared to those with just two polar cells, (mean migration was 3.8% (n=17) compared to 22.2% (n=36), respectively, t-test *P*=0.003), suggesting that larger *not^1^* clusters had particular difficulty in making their way successfully to the oocyte.

**Fig. 2.**
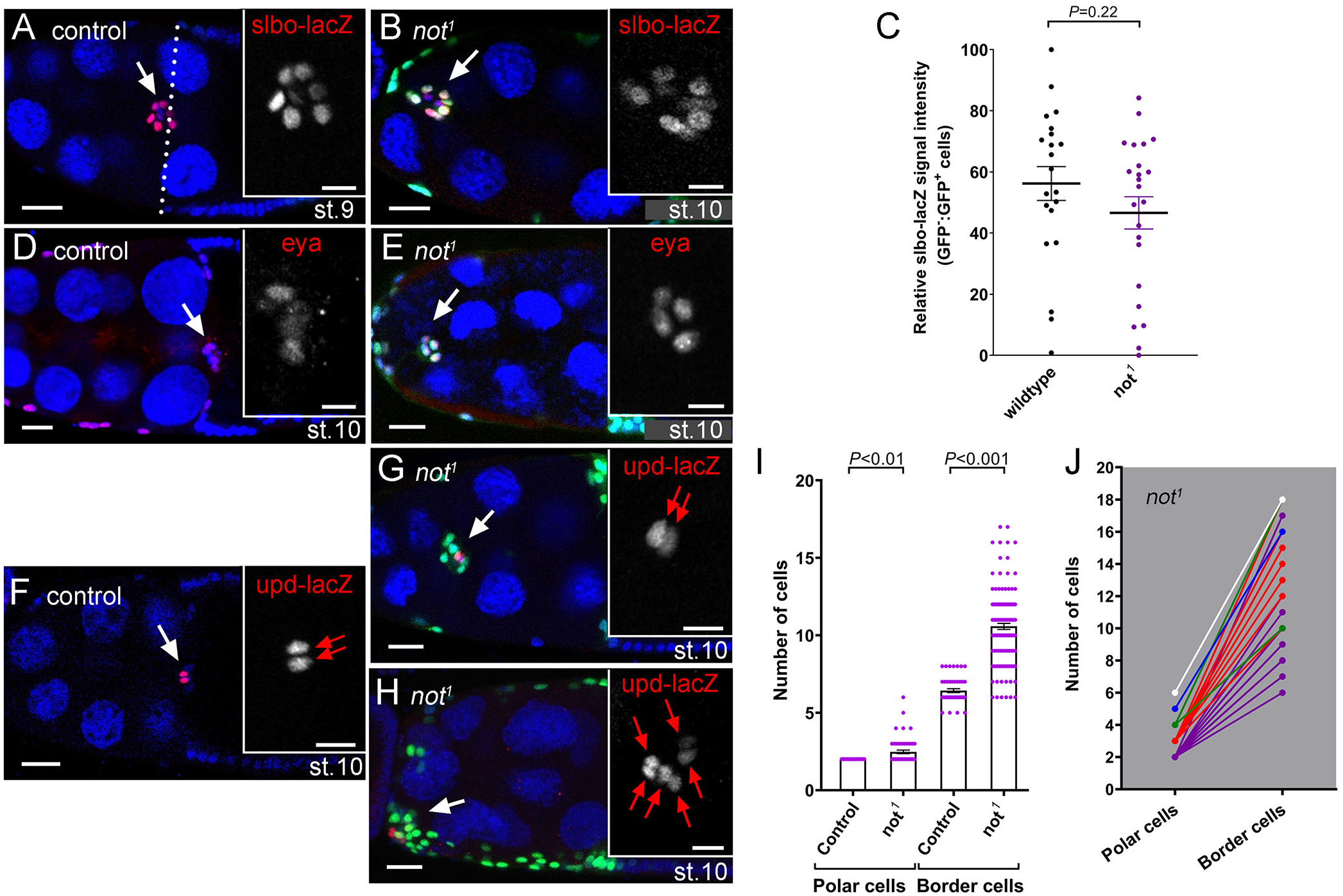
*non-stop* regulates polar cell number. **A**, Control egg chamber at mid stage 9 showing *slow border cells* expression with the *slbo-lacZ* reporter (red) in migrating border cells. Nuclei are labelled with TOPRO-3 (blue). Inset shows magnified image of *slbo-lacZ* alone (greyscale). **B**, Stage 10 egg chamber with *not^1^* mutant border cells labelled by MARCM with GFP (green). The normal pattern of *slbo-lacZ* is detected. **C**, Quantitation of relative *slbo-lacZ* signal intensity (GFP^−^, internal control: GFP^+^ homozygous sibling cell, see Methods for genotypes), showing no significant difference in *slbo* expression between wild type and *not^1^* mutant cells. **D**, Control egg chamber at stage 10 showing anti-Eyes absent antibody staining (Eya, red). Nuclei are labelled with TOPRO-3 (blue). Inset shows magnified image of Eya alone (greyscale). **E**, Stage 10 egg chamber with *not^1^* mutant border cells labelled by MARCM with GFP (green). In both control and *not^1^* mutant cluster cells Eya is restricted to outer border cells. **F**, Control egg chamber at stage 10 showing *unpaired* expression with the *upd-lacZ* reporter (red). Nuclei are labelled with TOPRO-3 (blue). Inset shows magnified image, showing *upd-lacZ* expression (greyscale) is restricted to the two polar cells (red arrows). **G-J**, 32% of *not^1^* mutant clusters possessed more than two *upd-lacZ* positive polar cells, which is associated with an increase in border cell numbers. **G**, GFP-labelled *not^1^* mutant border cell cluster possessing two *upd-lacZ^+^* nuclei, which represents the most abundant category, but some clusters contain up to 6 *upd-lacZ^+^* nuclei (**H**). **I**, Quantitation of polar and border cell numbers, reveals a significant increase in numbers of both *upd-lacZ^+^* polar cells and border cells in *not^1^* mutant clusters compared to controls. **J**, Graph showing the relationship between number of *upd-lacZ^+^* polar cells and border cells in individual *not^1^* border cell clusters, colour coded according to numbers of polar cells: 2, purple (n=36); 3, red (n=12); 4, green (n=3); 5, blue (n=1), 6, white (n=1). Bars in confocal images are 25 μm (10 μm for insets).

### *non-stop* is required for normal actin polarity in migratory border cells

Following their specification, border cells undergo two phases of cell migration, an initial polarised phase, and a second phase that utilises collective migration (Bianco et al., 2007). In the initial phase, leader cells exhibit long, highly polarised F-actin protrusions that are required for adhesion to and migration through the substratum (Fulga and Rorth, 2002). Later, F-actin accumulates around the cortex of the cluster, as cells alternate their position in the cluster as they move collectively (Bianco et al., 2007). In *not^1^* mutant clones, we observed a loss of initial F-actin polarity, and F-actin accumulation was subsequently not restrained to the cluster cortex but it also accumulated along border cell-border cell junctions (Fig. 3A,B). Quantification of F-actin staining confirmed a 2.6-fold shift in relative distribution towards the interior border cell junctions in *not^1^* mutant clusters compared to controls (two-way Anova P<0.0001, Fig.3C). This change in distribution was rescued by transgenic *not^+r^* overexpression (Fig.3C). When we examined egg chambers by live imaging, we found that progressive migration was reduced by 80%, from 0.45 μm/min in controls to 0.09 μm/min in *not^1^* mutant border cell clusters (P<0.01). This was accompanied by loss of initial F-actin polarity (Fig.4A,B, video S2 and S3) and a premature tumbling motion (Fig.4A-C). Further analysis revealed that while there was not a global reduction in the number of protrusions in *not^1^* mutant clusters (Fig.4D), there was a significant change in the distribution of the number (Fig.4E) and size (Fig.4F,G) of protrusions, from a front bias in controls (54% of protrusions) to the sides (63% of protrusions) in *not^1^* mutants (Fig.4E, P<0.01), consistent with a failure of these clusters to move in a polarised fashion.

**Fig. 3.**
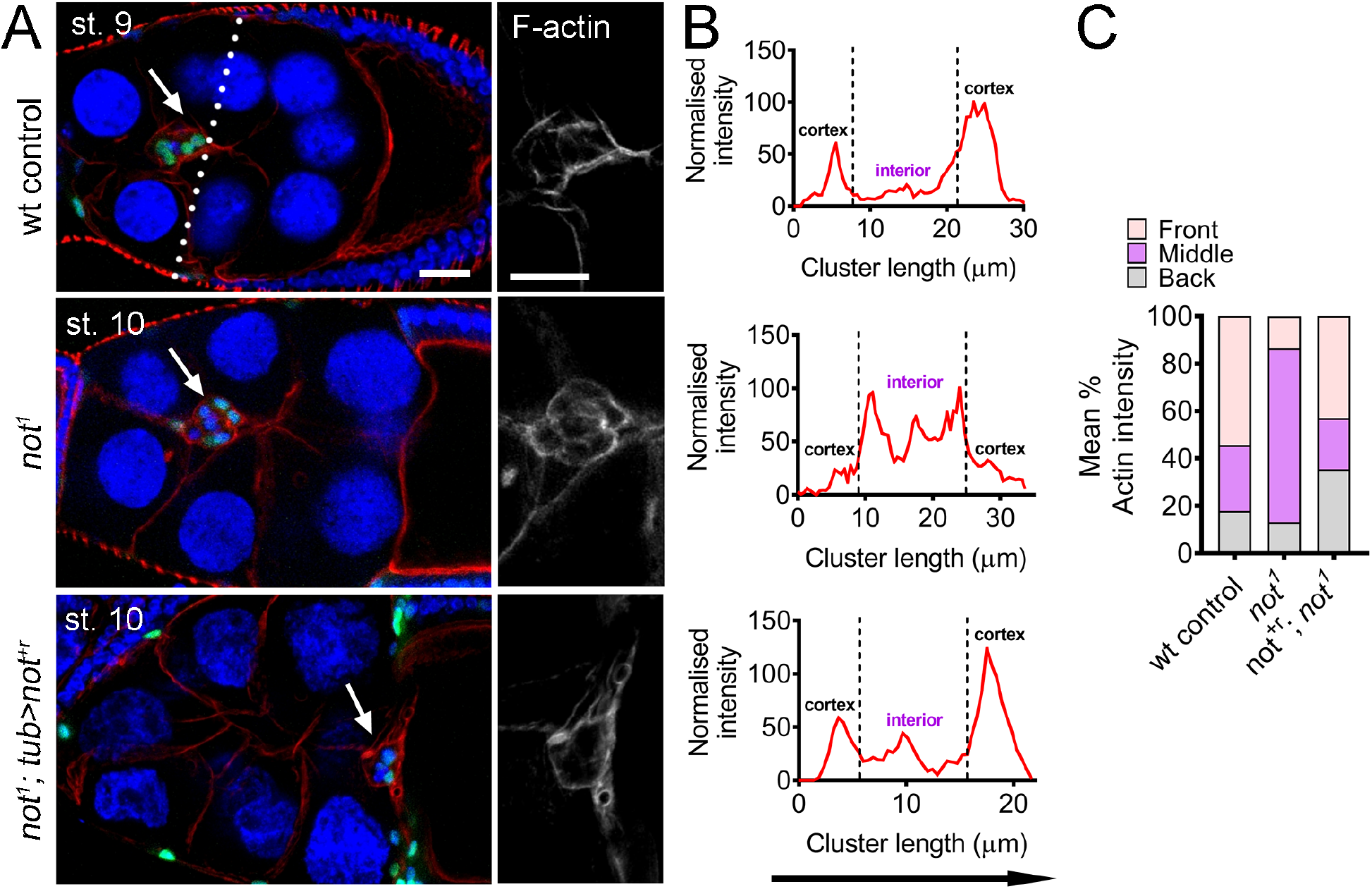
*non-stop* is required for normal actin polarity in migratory border cells. **A**, Confocal micrographs of egg chambers harbouring wild type, *not^1^* or rescued *not^1^* GFP-labelled clones (not^1^; tub>not^+r^) labelled with Phalloidin to visualise F-actin (red), TO-PRO-3 to label nuclei (blue). Egg chambers are stage 10 except the wt control, which is shown at mid-migration at stage 9 (dotted line indicates expected position of the cluster at this stage of migration). Border cell clusters are indicated with arrows. In wt, F-actin is normally polarised, with high levels around the cortex of the cluster, at border cell-nurse cell junctions. In contrast, in *not^1^* clusters, F-actin predominantly accumulates at internal border cell-border cell junctions; this is rescued by transgenic overexpression of *not^+r^*. Bars are 25 μm (RGB images) and 10 μm for magnified grayscale images of F-actin. **B**, Representative line scans of the same genotypes showing signal intensities of F-actin from anterior (left) to posterior (right), showing the change in F-actin profile in *not^1^* mutant clusters. **C**, Mean ratios of area under curve for front, middle and back of the cluster derived from lines scans taken from several egg chambers (wt control clusters, n=7; not^1^ clusters, n=9; tub>not^+r^, not^1^ clusters, n=16) showing a consistent defect in F-actin polarisation in *not^1^* clusters.

**Fig. 4.**
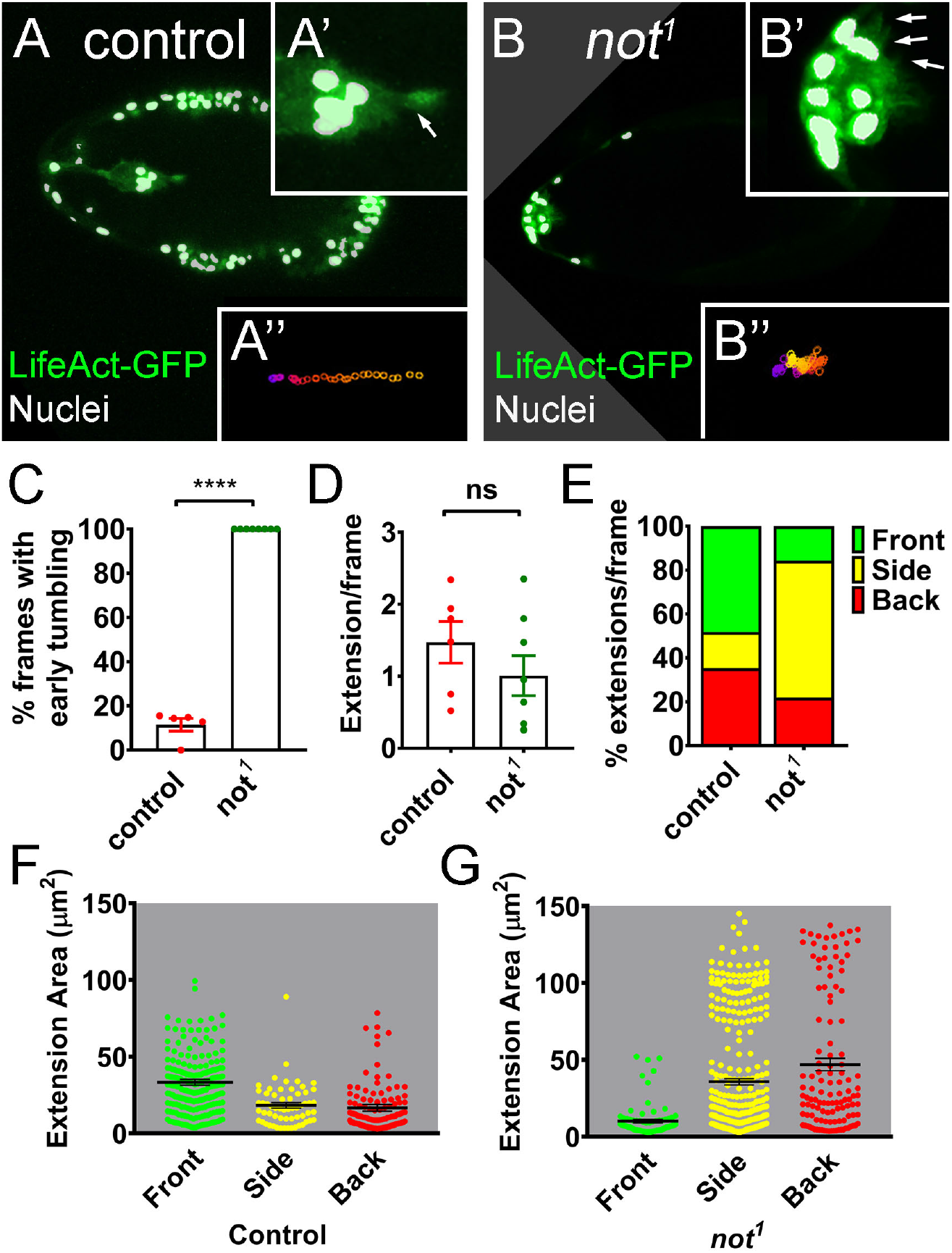
Loss of *non-stop* results in loss of polarised protrusions, retarded migration and early tumbling. **A-B**, Still images from time-lapse imaging of LifeAct-GFP-labelled border cells near the start of border cell migration with nuclear GFP-labelled MARCM clones labelled in white and LifeAct-GFP in green. **A** (inset **A’**, magnified image), Control egg chamber with clearly visible polarised F-actin protrusion at the leading edge of the cluster (arrow in **A’**), leading to progressive migration from anterior to posterior (track **A’’**, generated using a custom macro, (Poukkula et al., 2011)). In contrast, *not^1^* clusters (**B**) display multiple shorter protrusions at different positions around the cluster (arrows in **B’**), leading to poorly directed movement of the cluster towards the posterior pole (track **B’’**). **C-G**, Quantitation of time-lapse images from wt control (n=5) and *not^1^* (n=7) LifeAct-GFP-labelled border cell clusters, showing effects on tumbling and actin-based cellular protrusions. **C**, Graph showing percentage of frames from the first half of migration with tumbling border cells. Individual data points are shown together with mean ±SEM. *not^1^* significantly increases early tumbling ***, P<0.0001 Student’s t-test. **D**, Graph of total cellular extensions/frame after segmentation. There is no significant difference (ns, Student’s t-test) between wt control and *not^1^*. **E**, Graph of percentage extensions/ frame at front, back or sides of the cluster, showing a higher proportion of extensions at the side of *not^1^* clusters compared to controls. **F,G**, Measurements of the area of extensions detected at the front, back or sides of wt and *not^1^* clusters, together with mean area ±SEM, showing that the size of protrusions at the front is reduced in *not^1^* clusters concomitantly with an increase in the size of extensions at the side and back.

### *non-stop* acts independently of Scar during border cell migration

Recent data suggest Non-stop is capable of interacting with Arp2/3 and the WAVE regulatory complexes (WRC) in the cytoplasm to prevent polyubiquitination and subsequent proteasomal degradation of the WRC subunit Scar (Cloud et al., 2019). Scar/WAVE-Arp2/3 interactions result in nucleation of branched actin filament networks and in that way regulate migration (Buracco et al., 2019; Krause and Gautreau, 2014). This prompted us to test whether loss of *non-stop* function resulted in destabilisation of Scar levels in border cells. Endogenous Scar staining was very faint (Fig.5A,B) compared to ectopically overexpressed Scar (Fig.5C), but we did not observe any difference in Scar protein staining between *not^1^* mutant border cells and their heterozygous siblings (Fig.5A,B). To test whether *Scar* loss-of-function phenocopied *not^1^* clusters, we generated homozygous clones for an amorphic *Scar* mutant allele, *Scar^Δ37^* (Zallen et al., 2002). Notably, we found F-actin polarity was unaffected with F-actin being predominantly distributed at the cortex of *Scar^Δ37^* clusters. Migration of *Scar^Δ37^* clusters was retarded. However, previous live imaging analysis of clusters in which *Scar* had been knocked down by RNAi, revealed that *Scar* loss of function resulted in a reduction in the number of cellular protrusions, with a higher proportion of protrusions at the rear of the cluster, and fewer in the front and middle compared to controls (Law et al., 2013). These phenotypes are consistent with a reduction in migration, but not with the *not^1^* phenotypes described above (Fig.3,4). Polarisation of the polarity determinant Crb was also normal in *Scar* mutant clones suggesting the architecture of the clusters was unaffected. Taken together, we conclude that, in border cells, *non-stop* acts independently of *Scar* to drive collective migration.

**Fig. 5.**
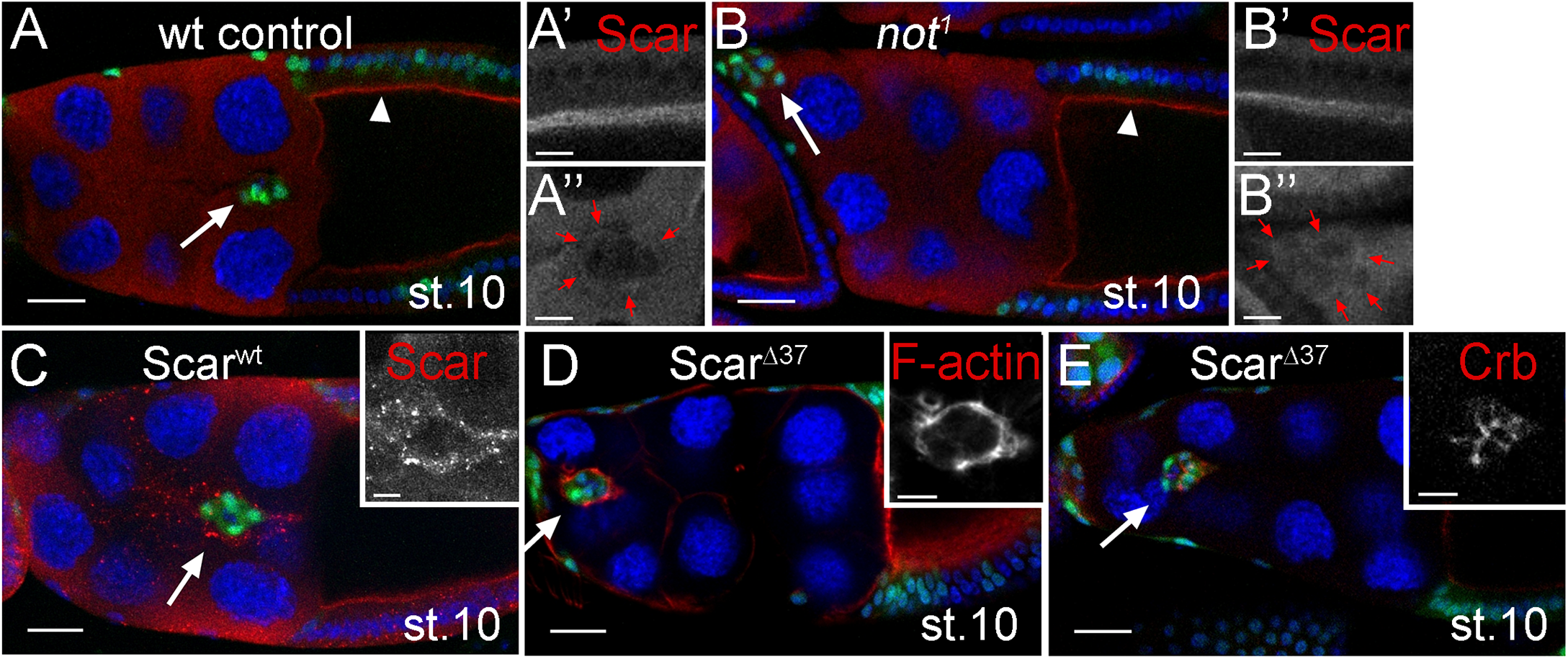
Non-stop acts independently of Scar during border cell migration. **A,B,** Scar levels are not reduced in *not^1^* follicle or border cells. Confocal micrographs showing Scar staining (red) in wt (**A**) and *not^1^* (**B**) GFP-labelled MARCM clones (green) at stage 10 of egg chamber development. Arrowhead, follicular epithelium; arrow, border cell. Bars, 25μm (magnified images, in greyscale, bar 10μm). **A’** (magnified image), Scar, shown in greyscale, predominantly localises to apical junctions of columnar follicle cells. **A’’** (magnified image), Scar staining is cytoplasmic in nurse cells but, in border cells, can be detected at the outer junctions of the cluster (red arrows). **B**, Scar is similarly localised in *not^1^* clones, with no reduction in level either at the apical side of follicle cells (**B’**) or at border cell junctions (**B’’**). **C**, Overexpression of wild type Scar (Scar^wt^ OE) using *slbo-GAL4* results in a robust signal confirming of Scar staining at the outer junctions of border cells. Arrow, border cells. Bars, 25μm (inset, 10μm). **D-E**, *Scar^Δ37^* clusters show normal F-actin and Crb polarity in migrating clusters. Shown are confocal micrographs of stage 10 egg chambers labelled with GFP (green) to mark clones of *Scar^Δ37^* cells induced with the MARCM technique and TOPRO-3 (blue) to stain all nuclei. Bars, 25μm (insets, 10μm). **D**, Egg chamber stained with Phalloidin showing localisation of F-actin around the cortex of the cluster (arrow); **E**, Egg chamber stained with antibodies against Crb, showing localisation to inner border cell junctions. *Scar^Δ37^* mutant border cell clusters display defective migration, with clusters frequently failing to reach the oocyte border by stage 10, as shown in these examples.

### *non-stop* is required for the normal level and/or distribution of Hippo signalling components in border cells

The loss of normal actin polarity, early tumbling of the border cell cluster, increased polar cell number are all features of Hippo signalling loss-of-function (Lin et al., 2014; Lucas et al., 2013). In outer border cells, the key upstream components of the Hippo pathway (Crumbs, Kibra, Expanded, Merlin) are found at sites of border cell-border cell contact (Lucas et al., 2013; Niewiadomska et al., 1999), where the pathway acts independently of the canonical downstream effector Yorkie to limit the activity, but not the recruitment, of the actin polymerisation protein Enabled (Lucas et al., 2013). This prompted us to test whether *non-stop* may be required for the normal level or distribution of Hippo signalling components in outer border cells. Using a transcriptional reporter of *expanded* expression (*ex-lacZ*), we found a 2.46 fold reduction in *expanded* levels in *not^1^* mutant cells compared to heterozygous sister cells in mosaic border cell clusters (Fig.6A,B; *P*=0.003, n=26). Similarly, we saw a reduction in Merlin protein levels at border cell-border cell junctions in *not^1^* mutant cells (Fig.6C,D). The distribution of Enabled appeared largely unaffected in *not^1^* mutant clusters (Fig.6E,F). In follicle cells, Expanded and Merlin are redundantly required for normal localisation of the apical transmembrane protein Crumbs (Crb) (Aguilar-Aragon et al., 2020; Fletcher et al., 2012). Strikingly, when we examined the distribution of Crb, we found that rather than being distributed in the junctions between neighbouring border cells (Niewiadomska et al., 1999), it was localised around the cortex of the cluster, at the interface between border cells and nurse cells (Fig.6G,H). Crb is required for polarisation of other polarity determinants, including aPKC in border cells (Wang et al., 2018). Correspondingly, the distribution of aPKC was somewhat disrupted in *not^1^* mutant cells (Fig.6I,J). We also observed a modest effect on the distribution of the adherens junction protein Armadillo/β-catenin (Arm; Fig.6K,L). Taken together, these data show that *non-stop* is required for expression of hippo signalling components and correct recruitment of polarity determinants in outer border cells.

**Fig. 6.**
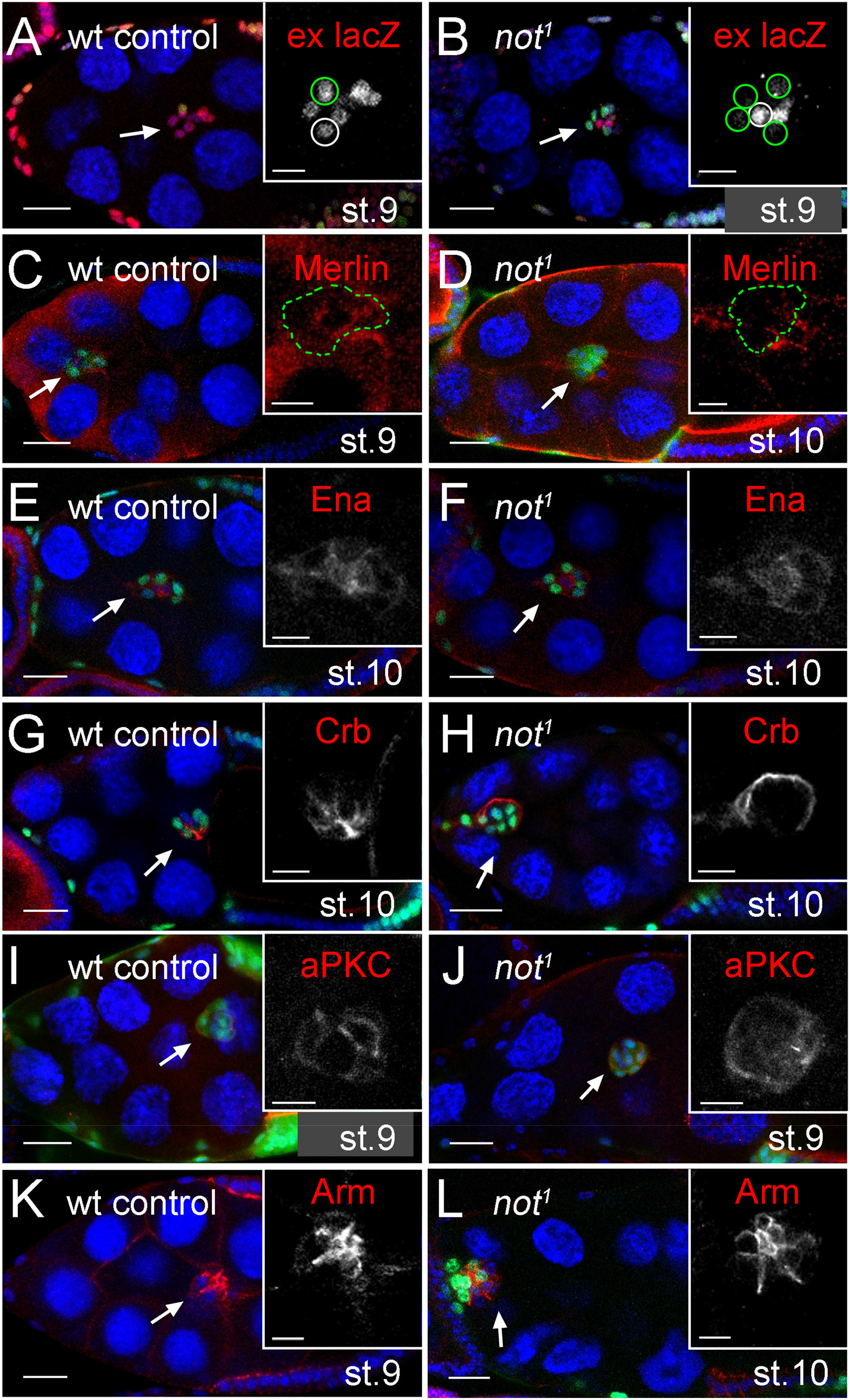
*non-stop* is required for the normal level and/or distribution of Hippo signalling components and polarity determinants in border cells. **A-L**, Confocal micrographs showing egg chambers with either wt or *not^1^* GFP-labelled MARCM clones (green) stained with antibodies against β-gal (to detect *ex-lacZ* expression, **A,B**); Merlin (**C,D**), Ena (**E,F**), Crb (**G,H**), aPKC (**I,J**), or Arm (**K,L**) in red. Nuclei are stained with TO-PRO-3 (blue). Bars 25 μm (10 μm for insets). Arrows, border cells. The stage of egg chamber development is as indicated. **A**, Mosaic order cell clusters, showing the normal expression of *ex-lacZ* in both GFP-labelled control clones (green outline), and control sibling cells (white outline). **B**, Notably, there is a reduction in *ex-lacZ* expression in *not^1^* clones (green outline) compared to control sibling cells (white outline). **C**, Merlin staining is weak but clearly detectable at the inner-border cell junctions in control clones (green outline), but **D**, is lost in GFP-labelled *not^1^* cells (green outline) and not adjacent control cells of the same cluster. **E-F**, Ena is predominantly located at cell junctions around the polar cells, and at inner and outer border cell membranes, in both control (**E**) and *not^1^* (**F**) clones. **G**, Crb is normally distributed at inner border cell junctions in control border cell clusters, but **H**, is strikingly redistributed to the cortex of *not^1^* border cell clusters. **I**, aPKC is normally distributed at inner border cell junctions in control border cell clusters, but **J**, this distribution is disrupted in *not^1^* clones, with some loss of aPKC at the inner membranes and a more cytoplasmic distribution in the border cells. **K**, the adherens junction protein Arm is apically localises at inner junctions in controls. **L**, in *not^1^* border cell clusters Arm appears more spread out, although remains restricted to inner junctions.

### *ex* and *mer* are targets of Non-stop but not the HAT module of SAGA, which is dispensable for border cell migration

A key and highly conserved role of Non-stop/ USP22 is to regulate gene expression, acting as a central component in the DUB module of the SAGA complex (Lee et al., 2011). By exploiting genome-wide ChIPSeq data from a recent study of the *Drosophila* SAGA complex (Li et al., 2017), we asked whether any of the canonical hippo signalling components are transcriptional targets of Non-stop. To do this we looked for binding sites at the gene promoters, −1000 to +200bp of the transcription start sites. We found that the *expanded* promoter is bound by Non-stop (n=2, Fig.7A); furthermore, depletion of *non-stop* leads to a 2.5 fold reduction *expanded* expression in embryos (Li et al., 2017), comparable to the effect we observed in border cells (see above). Similarly, we also found evidence that Not binds the *merlin* promoter (n=1, Fig.7B). Interestingly, Ada2b, a SAGA-specific HAT module subunit, that anchors the HAT module to SAGA and is required for its HAT activity (Kusch et al., 2003; Lee et al., 2011; Muratoglu et al., 2003; Pankotai et al., 2005; Zsindely et al., 2009), did not bind either of these loci (n=4, Fig.7A,B), suggesting that *expanded* and *merlin* promoters are DUB specific targets. Correspondingly, we did not see a reduction in *ex-lacZ* levels in *ada2b* mutant clones (Fig.7C). Furthermore, when we tested the requirement for *ada2b* in F-actin polarity and border cell migration, we found that *ada2b* mutant border cells migrated normally with cortically-localised F-actin (Fig.7D, mean migration 82.3% ±3.6%, n=45). Taken together, these data indicate that the DUB module can regulate transcription of *expanded* and *merlin* independently of the HAT module in border cells.

**Fig. 7.**
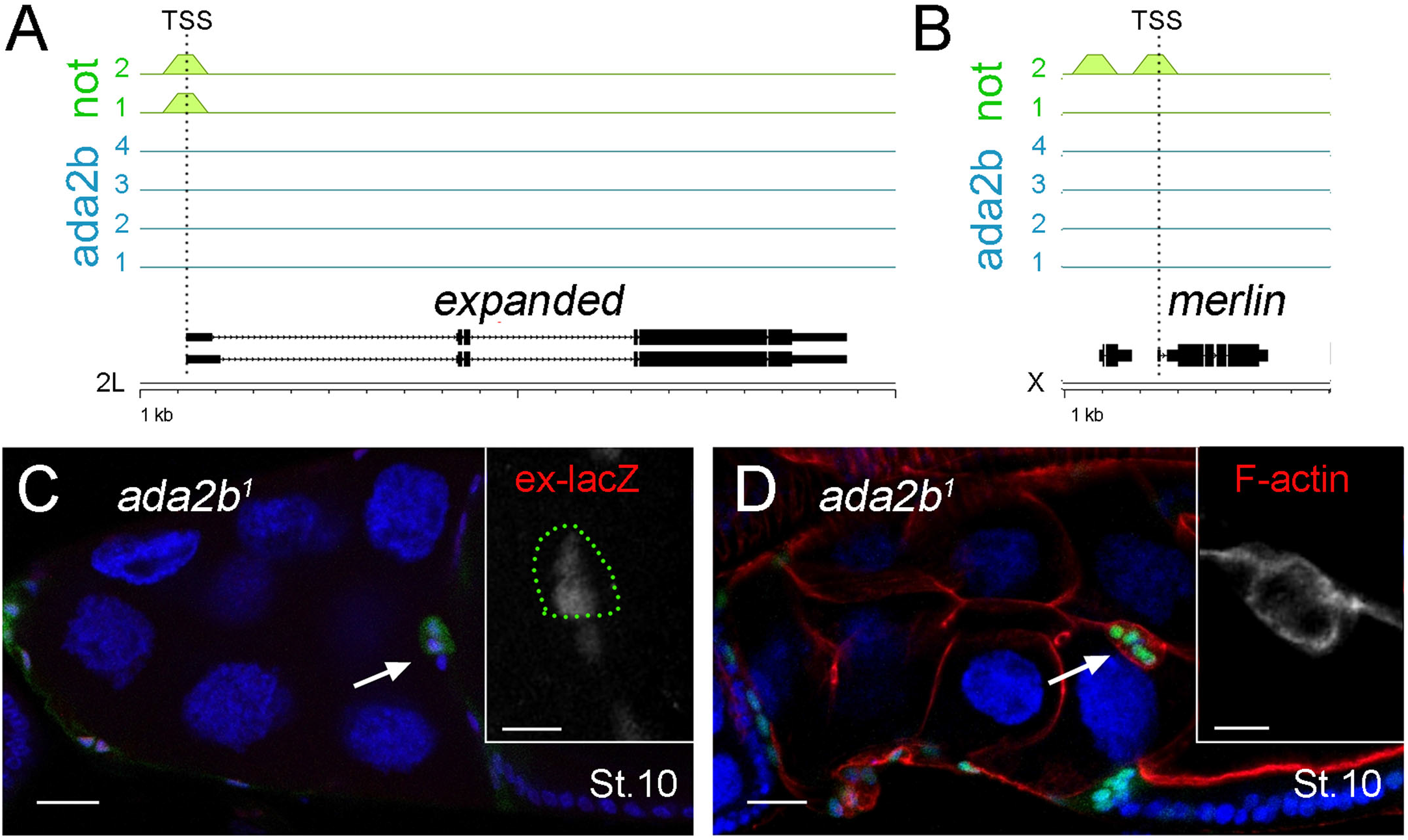
Ex and Mer are targets of Non-stop but not the HAT module of SAGA, which is dispensable for border cell migration. **A-B**, Non-stop, but not Ada2b bind to the *expanded* and *merlin* promoters. At the top are the ChIP binding profiles for all replicates of Not (green, n=2) and Ada2b (blue, n=4) at *expanded* and *merlin* promoters in *Drosophila* embryos as determined from data reported in (Li et al., 2017). Position of the transcription start site (TSS) is shown with a dotted line. Below is a schematic of the gene structure at the respective genomic loci with exons (thick lines) and introns (thin lines). Scale bar, 1 kb intervals. **C**, Confocal micrograph of a stage 10 egg chamber (arrow) with *ada2b^1^* GFP-labelled MARCM clone (green) stained with antibodies against β-gal (red) to detect *ex-lacZ* expression. Nuclei labelled with TO-PRO-3 (blue). Inset, *ex-lacZ* staining in grayscale, with mutant cells outlined (green dotted line). There is no reduction in *ex-lacZ* staining in *ada2b^1^* mutant cells compared to sibling control cells. **D**, Confocal micrograph of a stage 10 egg chamber (arrow) with *ada2b^1^* GFP-labelled MARCM clone (green) stained with Phalloidin to label F-actin (red), showing F-actin is localised to outer border cell junctions, as wild type, compare Fig2A.

### Overexpression of *ex* partially rescues cell migration and polarity defects

To further explore the functional significance of reduced *expanded* levels, we examined the effect of *expanded* loss-of-function on border cell polarity and migration (Fig.8). We found that border cells mutant for an *expanded* loss-of-function allele (*ex^e1^*) phenocopied the effect of *not^1^*, albeit more weakly (Fig.8A-F), with some loss of cortical F-actin staining and a significant disruption of Crumbs distribution (Fig.8K-L), accompanied by abrogated migration (Fig.8M). Strikingly, *expanded* overexpression (*ex^+^*) substantially restored more normal Crumbs and F-actin distributions in *not^1^* mutants (Fig.8G-H and K-L) and significantly suppressed the effect of *not^1^* on migration (Fig.8M; the mean percentage migration of *ex^+^ not^1^* border cell clusters was 55.2 ±3.0%, n=75 compared to 38.7 ±2.9%, n=101 for *not^1^* alone, P<0.0001). Taken together with the data above, we conclude that *expanded* is a critical transcriptional target of *non-stop* required for its function in border cells. Previous studies have shown that overexpression of Capping protein B (*cpb^+^*), which antagonises Enabled by competing for binding F-actin barbed ends and preventing actin polymerisation, is capable of complementing impaired hippo signalling (loss of *warts*) in border cells. Correspondingly, we find that *cpb^+^* has a similar ability as *ex^+^* to rescue *not^1^*-associated defects in F-actin polarity and collective cell migration (Fig.8I-M). Interestingly, we also saw a partial recovery in the Crb distribution in *cpb^+^ not^1^* border cell clusters (Fig.8K), indicating a role for the actin cytoskeleton in controlling Crb polarity.

**Fig. 8.**
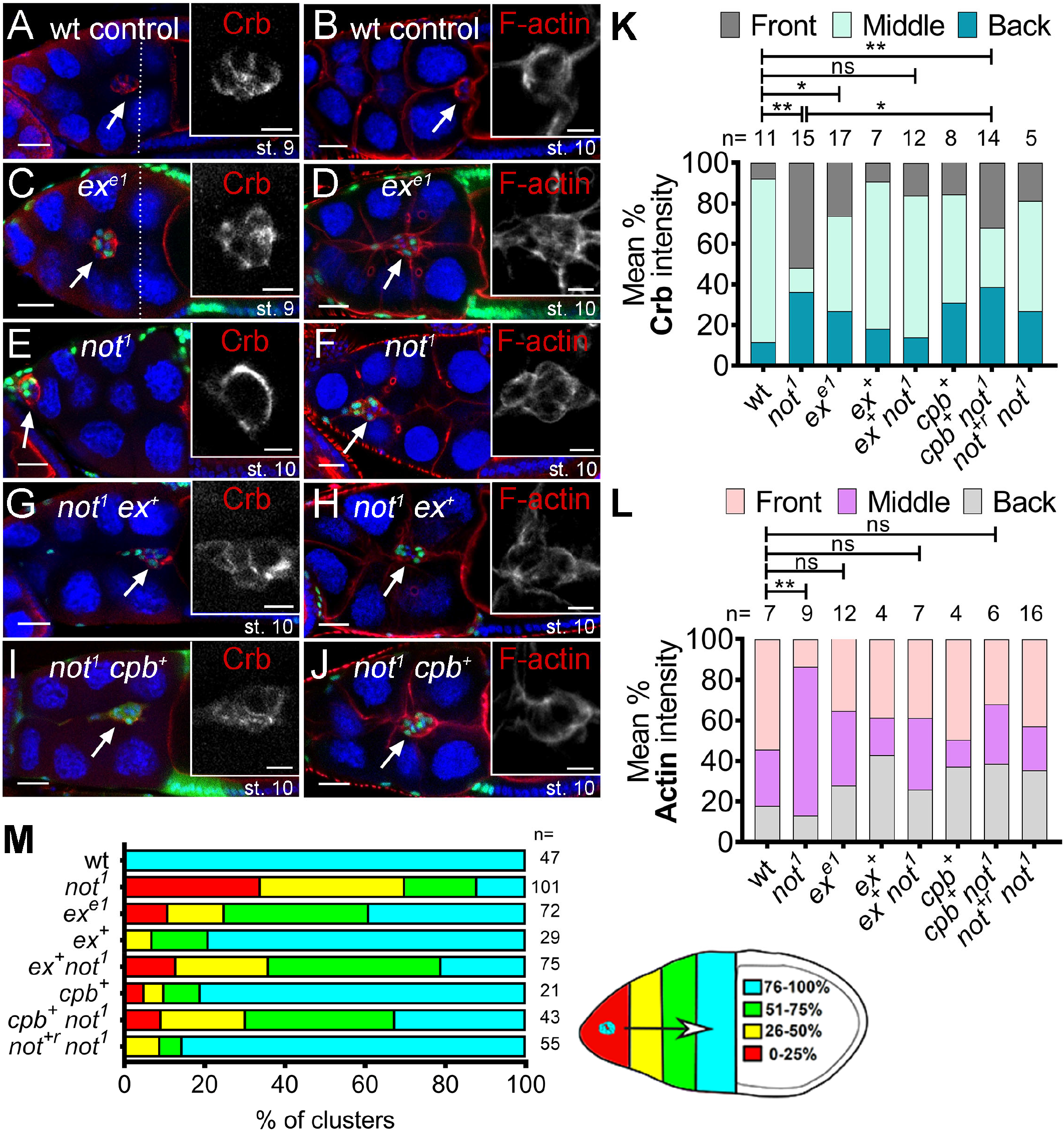
Overexpression of *expanded* or the actin capping protein *cpB* partially rescues cell migration and polarity defects. **A-J**, Confocal micrographs of Crb or F-actin (red) staining in egg chambers harbouring GFP-labelled MARCM clones (green) of different genotypes. TOPRO-3 (blue) labels all nuclei. Bars, 25μm (insets 10 μm). The stage of egg chamber development is as indicated, with dotted line showing position of overlying centripetal follicle cells in stage 9 chambers. Border cells are indicated with arrows. **A**, Control showing normal distribution of Crb at contacts between the border cells inside the cluster. **B**, Control showing normal cortical distribution of F-actin around the outer membrane of the cluster. **C**, *exe^1^* clones showing partial disruption of Crb. **D**, F-actin polarisation is also partially impaired in *exe^1^* border cells, with some F-actin visible at inner junctions of the migrating clusters. **E**, Crb is redistributed away from inner junctions to the cortex of the cluster in *not^1^* clones. **F**, F-actin is found distributed on inner junctions of *not^1^* clusters between border cells. **G**, The disruption of Crb localisation in *not^1^* clones is partially rescued by overexpression of *ex* (*not^1^ ex^+^*). **H**, F-actin also is more normally polarised in *not^1^ ex^+^* border cells, although some weak staining is also evident between border cell-border cell junctions. **I**, Overexpression of cpB weakly restores some Crb distribution in *not^1^* mutant cells (*not^1^ cpB^+^*) and **J**, F-actin is displaced from border cell junctions inside the *not^1^ cpB^+^* clusters. **K**, Quantification of mean percentage of Crb staining at the front, middle and back of the cluster (area under curve measurements) derived from lines scans taken from several egg chambers (n=number of clusters). * P<0.05; ** P<0.001; ns, not significant, 2-way Anova comparisons of mean ratio of Crb staining at the front of the cluster; comparable results were obtained for comparisons of staining in the middle of the cluster (not shown). **L**, Quantification of mean percentage of F-actin staining at the front, middle and back of the cluster (area under curve measurements) derived from lines scans taken from several egg chambers (n=number of clusters). ** P<0.001; ns, not significant, 2-way Anova comparisons of mean ratio of F-actin staining at the front of the cluster; comparable results were obtained for comparisons of staining in the middle of the cluster (not shown). **M**, Histograms summarising border cell migration defects at stage 10 in the indicated genotypes (n= number of egg chambers), alongside the migration index for quantification of migration.

## Discussion

### A *non-stop*-mediated transcriptional programme establishes F-actin polarity during collective migration

Here we report that *Drosophila* USP22, encoded by *non-stop*, is necessary for F-actin polarity and collective cell migration of invasive border cells. Collective border cell migration requires actomyosin polymerisation and contraction at the cortex around the cluster as it moves over the nurse cell substrate; F-actin is effectively excluded from the center of the cluster where polarity determinants acting via the Hippo complex block the activity of the F-actin regulator Enabled. Mechanistically, our experiments suggest *non-stop* regulates inside-out F-actin polarity by regulating the expression of hippo signalling components, *ex* and *mer*, which are direct Not targets. Not has been reported to regulate the actin cytoskeleton directly by promoting the stability of Scar/WAVE. However, we did not observe a reduction in Scar levels in *not* mutant clones and *Scar* loss-of-function did not disrupt F-actin polarity.

Furthermore, we did not observe a significant change in the number of actin protrusions following *not* loss of function, which might be expected if *Scar* were a target in border cells. Notably, we find that overexpression of *expanded* suppressed *not^1^*-induced F-actin accumulation at inner border cell junctions, consistent with partial restoration of Hippo function and inhibition of Enabled function. We also observed that *cpb* overexpression rescued loss of *non-stop*, again consistent with disruption of Enabled function due to competitive binding of Cpb to F-actin barbed ends and the inhibition of F-actin polymerisation at inner border cell junctions. Interestingly, studies of maternally-provided *not* in the early embryo have identified a requirement for *not* in membrane invagination and nuclear anchoring during cellularisation (Li et al., 2017). Invagination is driven by actin, which is highly polarised at the base of invaginating membranes, and transiently in apical microvilli. Enabled plays an important role in establishing actin dynamics during invagination (Grevengoed et al., 2003), raising the question of whether the regulatory network between Not and the Hippo complex we have uncovered also has a role to play in this context.

### *non-stop* regulates the distribution of polarity determinants

A striking effect of *not* loss of function in border cells is the redistribution of Crb from inner to outer border cell junctions. When we looked at possible effects of this on other polarity determinants, we found localisation of aPKC to the inside apical junction between border cells was disrupted, consistent with previous studies showing that Crb, acting together with the Par complex and endocytic recycling machinery, is necessary for ensuring its correct distribution (Wang et al., 2018). Mislocalised aPKC generates protrusions at the side and back of border cells (Wang et al., 2018), just as we have seen in *not^1^* clusters. Hence, whilst loss of Hippo components leads to loss of inside-out actin polarity, disruption of Crb and aPKC might account for the change in orientation of protrusions. Why is Crumbs mis-localised to the cortex of the border cell complex? Our complementation experiments (Fig. 8) suggest that this might be at least partially accounted for by loss of expression of the FERM domain proteins Expanded and Merlin, which in follicle cells act together with Moesin (Moe) to recruit Crb to the apical surface (Aguilar-Aragon et al., 2020). Moe stabilises Crb at the apical membrane of epithelia by linking Crb to cortical actin (Medina et al., 2002). Although the physical interaction between Moe and Crb may be weak (Sherrard and Fehon, 2015), Moe is an important regulator of dynamic Crb localisation in follicle cells, as it acts to antagonise interactions between Crb and aPKC at the marginal zone of the apical membrane domain, while stabilising interactions between Crb and the apical surface (Sherrard and Fehon, 2015). Importantly, in border cells, Moe is cortically localised where it organises a supercellular actin cytoskeleton network and promotes cortical stiffness (Ramel et al., 2013). An attractive hypothesis therefore is that Moe, perhaps along with other proteins, is a sink for Crb at the cortex of the border cell cluster following loss of Ex and Mer at inner border cell junctions in *non-stop* mutants. When we overexpressed *expanded* the normal pattern of Crb localisation was partially restored, in support of a competitive binding model. Interestingly, we also observed weak rescue of Crb localisation following CpB overexpression. This might be because Moe, or other proteins that tether Crb on the outer membrane are only accessible in the absence of a strong supercellular F-actin cortex and that restoration of cortical F-actin in *not^1^ cpb^+^* cells displaces Crb. In wild type border cells, Crb needs to be constantly moved from the outside membrane in a dynamin- and Rab5-dependent manner (Wang et al., 2018). Another possibility therefore, which is not mutually exclusive with the first, is that polarisation of the F-actin cytoskeleton is important for correct trafficking of Crb in border cells, as it is in follicle cells (Aguilar-Aragon et al., 2020).

### *non-stop* is necessary for the expression of *ex* and *mer* independently of the requirement for *yki*

Abnormal accumulation of F-actin in epithelial tissues, e.g. resulting from loss of CpB, has been shown to lead to Yki-induced expression of *ex*, *mer*, and other target genes to reinforce Hippo activity at the cortex (Fernandez et al., 2011; Ko et al., 2016; Sansores-Garcia et al., 2011). It is known that the Hippo pathway integrates multiple inputs at the level of Yki and that Yki interacts with a number of chromatin-modifying factors for transcriptional activation of target genes (Hillmer and Link, 2019). Is it possible that *non-stop* acts to support Yki-mediated expression of *ex* and *mer*? In border cells, ectopic overexpression of Yki has been reported to accelerate border cell migration, resulting in clusters prematurely reaching the oocyte during stage 9, suggesting that there may be a Yorkie-mediated negative feedback loop to maintain F-actin homeostasis (Lucas et al., 2013). However, *yki* mutant border cells or clusters in which *yki* has been knocked down in the outer border cells migrate normally, suggesting that *yki* is normally dispensable in outer border cells for invasive migration (Lin et al., 2014). We therefore favour a model whereby *non-stop* provides independent transcriptional control of *ex* and *mer* in this context. The situation is different in polar cells, where the Hippo pathway is proposed to act by suppressing Yki activity and cell proliferation to maintain normal polar cell numbers. However, similar to Hippo loss-of-function or *yki* gain-of-function, we find that loss of *non-stop* leads to increased numbers of polar cells, which, again, argues against a role for *non-stop* in supporting *yki*-mediated gene expression. Nevertheless, what this does suggest is that the requirement for *non-stop* in Hippo complex formation is not limited to situations where the Hippo complex acts in a *yki*-independent fashion.

### SAGA-independent roles for *non-stop* during development and disease

The growth, specification and migration of cells during tissue development requires precisely regulated patterns of gene expression, that depend on numerous cues for temporal and spatial gene activation, involving crosstalk with multiple signalling pathways. Strikingly, it has emerged that factors once considered to be ubiquitous regulators of transcription, including the SAGA chromatin-modifying complex, can have specific roles in discrete developmental processes. Although it has been suggested that SAGA is required for all transcribed genes in some contexts (Bonnet et al., 2014), numerous studies have shown that loss of SAGA components affects the expression of only a subset of genes (Pahi et al., 2015; Pankotai et al., 2013; Zsindely et al., 2009) and different components modulate distinct and overlapping subsets (Helmlinger et al., 2008; Helmlinger et al., 2011; Lee et al., 2000; Weake et al., 2008). These differences in expression are likely to explain their different physiological roles; for instance, during female germline development in *Drosophila*, *ada2B* affects the expression of many genes and is required for oogenesis, whereas *non-stop* affects relatively few and is dispensable (Li et al., 2017). Elegant genome-wide ChIP studies indicate that even though both DUB and HAT modules bind the same genes, many of the targets do not require the DUB module for expression, explaining the observed dependencies. These experiments also reveal non-overlapping sites of chromatin occupancy for the DUB and HAT modules of SAGA in *Drosophila* (Li et al., 2017), but the significance of differences in transcriptional targeting for cell function had not been established. Notably, in this respect, we find that the requirement for *non-stop* in border cell migration is not matched by a requirement for *ada2b*. Furthermore, Ada2b has not been found to bind the *ex* and *mer* promoters, providing a molecular explanation for *non-stop*’s SAGA-independent role. Importantly, these findings challenge the perceived view that transcriptional roles for non-stop/USP22 are mediated solely by SAGA. This may have broader relevance to situations where USP22, but not other members of SAGA are associated with human disease states, particularly where cell polarity is frequently disrupted, such as cancer (Glinsky et al., 2005). Our current efforts are directed at identifying SAGA-independent factors that facilitate non-stop’s chromatin binding and function.

## Methods

### Non-stop transgene

An RNAi-resistant, full-length *non-stop* expression construct was synthesised by GeneArt (Invitrogen). RNAi-resistance was achieved by incorporating numerous silent polymorphic mutations, such that, in the regions targeted by dsRNAs, homology with the inverted repeat sequences was limited to no more than 8 contiguous base pairs (Jonchere and Bennett, 2013). The *non-stop* open reading frame was shuttled into pPMW-attB (Chen et al., 2015) by gateway cloning, placing the *non-stop* open-reading frame downstream of a Myc epitope tag. Stable transgenic flies were made by phiC31 integrase-mediated transgenesis at a landing site on the second (attP40, at 25C6) and third (attP2, at 68A4) chromosomes by the Cambridge fly facility (University of Cambridge).

### Drosophila stocks and genetics

Flies were raised and crossed at 25°C according to standard procedures. *w^1118^* or *FRT80B* flies were used as the wild-type control strains. 138 RNAi lines, corresponding to 45 *Drosophila* DUBs (details available on request), were screened for border cell defects at 25°C. *UAS-not^IR^* (Vienna Drosophila Resource Center #45776) was identified as having the most severe effect on migration. The FLP/FRT site-specific recombination system was used to generate mutant clones with a heat-shock promoter (Xu and Rubin, 1993). The following fly lines were obtained from the Bloomington Drosophila Stock Center: *FRT80B (*BL1988), *w^1118^* (BL6409), *slbo-Gal4, UAS-GFP* (BL6458, Montell Lab), *slbo-lacZ* enhancer trap line (BL12227), *slbo-Lifeact-GFP* (BL58364), c*306-Gal4, UAS-GFP* (BL3743). For clonal analysis we used the following strains:

*hsFLP, tub-Gal4, UAS-GFP; +/+; tubGAL80 FRT80B/TM6B* (generated from BL42732, BL5191),

*hsFLP, tub-Gal4, UAS-GFP; tubGAL80 FRT40A/+; +/TM6B* (generated from BL42732, BL5192),

*hsFLP, tub-Gal4, UAS-GFP; +/+; FRT82B tubGAL80/TM6B* (generated from BL42732, BL44408).

The amorphic *not* allele, *not^1^* was obtained from Margarete Heck and recombined with *FRT80B*. *FRT82B Ada2B* was a gift from Jerry Workman (Li et al., 2017). *UAS-Scar* and *FRT40A Scar^Δ37^* were gifts from Eyal Schejter. *UAS-cpB, UAS-ex* (Lucas et al., 2013), *upd-lacZ* (Jiang et al., 2009) and *ex-lacZ* (Fletcher et al., 2012), were gifts from Nic Tapon. Information on these strains is also available at http://www.flybase.org.

### Generation of mosaic clones using MARCM

Mosaic Analysis with a Repressible Cell Marker (MARCM) was used to generate positively marked clones labelled with GFP (Lee and Luo, 2001). Expression of genes under GAL4-UAS is inhibited in the presence of GAL80. Heat shocking induces the expression of heat shock (hs) driven FLP, which acts to induce recombination at Flippase Recognition Targets (FRT). Homozygous daughter cells lacking GAL80 are then capable of GAL4-mediated gene expression of GFP and other UAS-transgenes. Mitotic recombination is initiated after heat shock where some daughter cells are GFP^+^ while others are GFP^−^ due to the presence of GAL80. To obtain border cell mitotic (mosaic) clones, progeny of the right genotypes were heat shocked twice a day for 1 hour each with at least 5 hr intervals between treatments, from pupae to adult at 37°C. Newly enclosed adults (2-3 d old) were fattened for 2 d on yeast paste.

### Immunofluorescent staining

Ovaries were dissected in PBS (Phosphate buffer saline) and fixed with 3.7% paraformaldehyde in PBS. The ovaries were washed with PBST (1x PBS, 0.2% Tween 20) 3 times for 15 minutes each time. Ovaries were then blocked with PBTB (1x PBS, 0.2% Tween 20, 5% fetal bovine serum) for 1 hour at room temperature. The ovaries were treated with primary antibodies in PBTB at 4°C overnight. The following primary antibodies were used for immunostaining. developmental Studies Hybridoma Bank (DSHB): mouse anti-Armadillo (N27A1, 1:200, concentrate), mouse anti-Enabled (5G2, 1:25, concentrate), mouse anti-β-gal (40-1a, 1:300, concentrate), mouse anti-eyes absent (eya10H6, 1:100, supernatant), mouse anti-SCAR (P1C1, 1:200, concentrate). Mouse anti-aPKC ζ (sc-17781, 1:200) from Santa Cruz. Guinea pig anti-Merlin (1:7500) from R Fehon lab. The primary antibodies were washed with PBST 3 times 15 min and then blocked with PBTB for 1 hr at room temperature. Ovaries were incubated with Alexafluor-conjugated secondary antibodies (1:500, Life technologies) in PBTB at 4°C overnight. Phalloidin 555 (1:50, Molecular Probes) was used to stain F-actin. Ovaries were washed with PBST for 15 minutes before staining nuclei with TO-PRO-3 (Life technologies, 1:1000) in PBST for 15 minutes. Ovaries were mounted in Vectashield (Vector laboratories). For Crumbs staining, Ovaries were dissected in PBS (Phosphate buffer saline) and fixed with boiled 8% paraformaldehyde in PBS and heptane (6:1) for 10 minutes. Samples were treated with heptane and methanol (1:2) for 30 seconds. They were then washed in methanol for 10 minutes. The ovaries were washed with PBST (1x PBS, 0.2% Tween 20) 2 times for 15 minutes each time. Ovaries were then blocked with PBTB (1x PBS, 0.2% Tween 20, 5% fetal bovine serum) for 30 minutes at room temperature. The ovaries were treated with mouse anti-Crumbs (Cq4, 1:100, concentrate, DSHB) in PBTB at 4°C overnight.

### Image acquisition and analysis of fixed samples

Images were taken on a confocal microscope (LSM710 or LSM780, Carl Zeiss) using 20x/0.5NA air objectives. Three laser lines were used based on the excitation of wavelength of the staining dyes which includes 488 nm, 561 nm and 633 nm wavelengths. Extent of migration (the migration index) was measured as a percentage of the distance travelled to the oocyte/nurse cell boundary in stage 10 egg chambers. ImageJ (https://imagej.nih.gov/ij/) was used for quantification of signal intensities in mosaic clusters using z-stack maximum projections. Raw integrated density was used as intensity values. For line scan profiles, maximum intensity images of Actin and Crumbs staining were generated in ImageJ. Background signal were subtracted. The plot profile function in Image J was used to measure signal intensities along lines drawn through the centre of border cell clusters and the peak analyser tool in OriginPro (Origin Lab) was used to calculate the area under peaks that were identified. The ratio of intensities at front, middle and back, were compared and normalised in Prism8 (Graphpad). The following statistical tests were performed using Prism 8 (GraphPad): Student’s t-tests; one-way or two-way Anova, with Tukey correction for multiple comparisons; multiple linear regression with least squares. Figures were made using FigureApp in OMERO (Allan et al., 2012; Burel et al., 2015) and final assembly in Adobe Photoshop.

### Egg chamber culture and time-lapse imaging of live egg chambers

Live imaging of egg chamber culture were as previously described (Law et al., 2013; Prasad et al., 2007) with slight modification. Briefly, media for both dissection and live-imaging, comprised of Schneider media (Gibco), 15% fetal bovine serum, 0.1 mg/ml acidified insulin (Sigma), 9 μM FM4-64 dye (Molecular Probes) and 0.1 mg/ml Pen-strep (Gibco) was freshly prepared. The pH of the media was adjusted to 6.90-6.95. Individual egg chambers from well fattened progeny of the right genotype were dissected and transferred to borosilicate glass bottom chambered coverglasses (ThermoFisher) for imaging. Imaging was done at 25°C. Time-lapse movies were acquired on an inverted confocal microscope (LSM 710; Carl Zeiss) using 20x/0.5NA air objectives. Two laser lines were used based on the excitation of wavelength of the endogenous GFP and FM4-64 dye, which are 488 nm, and 561 nm wavelengths respectively. 16-20 slices of Z-stacks were taken with 2.5 μm slices every 3 min.

### Analysis of time-lapse images

Time-lapse image analyses were performed using a custom macro for ImageJ to analyse the behaviour of border cell migration and extension dynamics (Law et al., 2013; Poukkula et al., 2011) with slight modification. Briefly, time-lapse movies were split into different channels. Maximum projections of the GFP-channel were created. Egg chambers were rotated so that anterior ends were at the left. Border cells were manually thresholded to mask nuclear GFP generated from the MARCM system through the first or early phase of migration. Images of border cells clusters were then segmented into cell body and cellular extensions using signals from *slbo-LifeAct-GFP*. Extensions were grouped based on their positions in relation to the leading edge of the cluster: front (315-45°), side (45-135° or 225-315°) and back (135-225°). The macro also enabled tracking of the movement of cluster to measure the migration speed. Forward directed speed was calculated on x-axis by taking distance of the centre of cluster at one time point relative to the next time point. The tumbling index was calculated as the mean percentage of frames per time lapse movie that showed rounded clusters, exhibiting changes in the position of individual cells within the cluster for two or more consecutive frames in the first half of migration. Data were collated in Microsoft Excel and independent Student’s t-tests were done with Prism 8 (GraphPad). For visualisation of stills (Fig4A,A’-B,B’), GFP-labelled nuclei were segmented in Imaris (Bitplane) and labelled in white.

### Analysis of previously reported ChIP datasets

ChIP-seq data were downloaded from GEO (https://www.ncbi.nlm.nih.gov/geo/) using accession GSE98862; the dm3 assembly of the *D. melanogaster* genome was obtained from UCSC (http://www.genome.ucsc.edu/cgi-bin/hgTables). Peaks from Ada2b and Non-stop ChIP experiments were mapped to the dm3 genome assembly using BEDtools software (Quinlan and Hall, 2010), and any genes matching to peaks from −1000 to +200 of the transcription start site (TSS) were identified. For visualisation of ChIP-seq peaks on the genome, we utilised the ‘karyoploteR’ R/Bioconductor package (Gel and Serra, 2017).

## Genotypes of strains

**Fig 1.**

A. *w^1118^/+; Slbo-Gal4, UAS-GFP/+*

B. *Slbo-Gal4, UAS-GFP/UAS-not^IR^*

C. *c306-Gal4, UAS-GFP; UAS-not^+r^/+*

E. (as A-C with) *Slbo-Gal4*, UAS-GFP/UAS-not^IR^; UAS-not^+r^/+

F. *hsFLP, tub-Gal4, UAS-GFP/+ ;; +, FRT80B/tub-Gal80, FRT80B*

G,I. *hsFLP, tub-Gal4, UAS-GFP/+ ;; not^1^, FRT80B/tub-Gal80, FRT80B*

H. (as F,G with) *hsFLP, tub-Gal4, UAS-GFP/+ ; UAS- not^+r^/+ ; not^1^, FRT80B/tub-Gal80, FRT80B*

**Fig. 2**

A. *hsFLP, tub-Gal4, UAS-GFP/+; slbo-lacZ/+; +, FRT80B/tub-Gal80, FRT80B*

B. *hsFLP, tub-Gal4, UAS-GFP/+; slbo-lacZ/+; not^1^, FRT80B/tub-Gal80, FRT80B*

C. Quantification of A,B:

<u>Wild type GFP^−^</u>: *hsFLP, tub-Gal4, UAS-GFP/+; slbo-lacZ/+; +, FRT80B/tub-Gal80, FRT80B* (or homozygous for *tub-Gal80, FRT80B*)

<u>Wild type GFP^+^</u>: *hsFLP, tub-Gal4, UAS-GFP/+; slbo-lacZ/+; +, FRT80B/+, FRT80B*

<u>*not^1^* GFP^−^</u>: *hsFLP, tub-Gal4, UAS-GFP/+; slbo-lacZ/+; not^1^, FRT80B/tub-Gal80, FRT80B* (or homozygous for *tub-Gal80, FRT80B*)

<u>*not^1^* GFP^+^</u>: hsFLP, tub-Gal4, UAS-GFP/+; slbo-lacZ/+; not^1^, FRT80B/ not^1^, FRT80B

D. *hsFLP, tub-Gal4, UAS-GFP/+ ;; +, FRT80B/tub-Gal80, FRT80B*

E. *hsFLP, tub-Gal4, UAS-GFP/+ ;; not^1^, FRT80B/tub-Gal80, FRT80B*

F. *hsFLP, tub-Gal4, UAS-GFP/upd-lacZ ;; +, FRT80B/tub-Gal80, FRT80B*

G,H. *hsFLP, tub-Gal4, UAS-GFP/upd-lacZ ;; not^1^, FRT80B/tub-Gal80, FRT80B*

I,J. (quantification of F-H)

**Fig. 3**

*wt* control: *hsFLP, tub-Gal4, UAS-GFP/+ ;; +, FRT80B/ tub-Gal80, FRT80B*

*not^1^: hsFLP, tub-Gal4, UAS-GFP/+ ;; not^1^, FRT80B/tub-Gal80, FRT80B*

*not^1^; tub>not^+r^: hsFLP, tub-Gal4, UAS-GFP/+ ; UAS-not^+r^/+ ; not^1^, FRT80B/tub-Gal80, FRT80B*

**Fig. 4**

Control: *hsFLP, tub-Gal4, UAS-GFP/+; slbo-LifeAct-GFP/+; +, FRT80B/tub-Gal80, FRT80B*

*not^1^: hsFLP, tub-Gal4, UAS-GFP/+; slbo-LifeAct-GFP/+; not^1^, FRT80B/tub-Gal80, FRT80B*

**Fig. 5**

A, A’ and A’’. *hsFLP, tub-Gal4, UAS-GFP/+ ;; +, FRT80B/tub-Gal80, FRT80B*

B, B’ and B’’. *hsFLP, tub-Gal4, UAS-GFP/+ ;; not^1^, FRT80B/tub-Gal80, FRT80B*

C. *Slbo-Gal4, UAS-GFP/UAS-Scar^wt^*

D-E. *hsFLP; tubGAL80, FRT40A/ Scar^Δ37^, FRT40A; Act>CD2>Gal4, UAS-GFP/+*

**Fig 6.**

A (wt control): *hsFLP, tub-Gal4, UAS-GFP/+ ; ex-lacZ/+; +, FRT80B/tub-Gal80, FRT80B*

B (*not^1^*): *hsFLP, tub-Gal4, UAS-GFP/+ ; ex-lacZ/+; not^1^, FRT80B/tub-Gal80, FRT80B*

C, E, G, I, K (wt control): *hsFLP, tub-Gal4, UAS-GFP/+ ;; +, FRT80B/tub-Gal80, FRT80B*

D, F, H, J, L (*not^1^*): *hsFLP, tub-Gal4, UAS-GFP/+ ;; not^1^, FRT80B/tub-Gal80, FRT80B*

**Fig. 7**

C. *hsFLP, tub-Gal4, UAS-GFP/+ ; ex-lacZ/+; ada2b^1^, FRT82B/tub-Gal80, FRT82B*

D. *hsFLP, tub-Gal4, UAS-GFP/+ ;; ada2b^1^, FRT82B/tub-Gal80, FRT82B*

**Fig. 8**

A, B. *hsFLP, tub-Gal4, UAS-GFP/+; ; +, FRT80B/tub-Gal80, FRT80B*

C, D. *hsFLP; ex^e1^, FRT40A/tub-Gal80, FRT40A; Act>CD2>Gal4, UAS-GFP/+*

E, F. *hsFLP, tub-Gal4, UAS-GFP/+; ; not^1^, FRT80B/tub-Gal80, FRT80B*

G, H. *hsFLP, tub-Gal4, UAS-GFP/+; UAS-ex^+^/+; not^1^, FRT80B/tub-Gal80, FRT80B*

I, J.*hsFLP, tub-Gal4, UAS-GFP/+; UAS-cpB^+^/+; not^1^, FRT80B/tub-Gal80, FRT80B*

K-M, (quantitation of A-J together with the following genotypes)

*hsFLP, tub-Gal4, UAS-GFP/+; UAS-ex^+^/+; +, FRT80B/tub-Gal80, FRT80B*

*hsFLP, tub-Gal4, UAS-GFP/+; UAS-cpB^+^/+; +, FRT80B/tub-Gal80, FRT80B*

*hsFLP, tub-Gal4, UAS-GFP/+; UAS-not^+r^/+ ; not^1^, FRT80B/tub-Gal80, FRT80B*

## Online supplementary material

**Video S1**: 4 h time-lapse of border cell migration starting from specification of the cluster and the ability of the cluster to acquire forward protrusion, followed by cell-on-cell migration to the anterior border of the oocyte. GFP expression is driven by *slbo-Gal4* to label the border cell cluster in green. Nuclei are labelled with *Ub-His2A-RFP* in magenta.

**Video S2**: 4 h time-lapse movie of normal border cell migration showing onset of migration including the ability of cluster to acquire forward actin protrusions. MARCM clones are labelled with nuclear GFP, F-actin is labelled with LifeAct-GFP. Egg chamber genotype: *hsFLP, tub-Gal4, UAS-GFP/+; slbo-LifeAct-GFP/+; +, FRT80B/tub-Gal80, FRT80B.*

**Video S3**: 4 h time-lapse movie of abnormal border cell migration showing early tumbling of the cluster and multi-directional actin protrusions in *not^1^* mutant cells labelled with nuclear GFP using MARCM. F-actin is labelled with LifeAct-GFP. Egg chamber genotype: *hsFLP, tub-Gal4, UAS-GFP/+; slbo-LifeAct-GFP/+; not^1^, FRT80B/tub-Gal80, FRT80B*

## Acknowledgments

We thank Rick Fehon, Margarete Heck, Timothy Megraw, Eyal Schejter, Nic Tapon, Jerry Workman the developmental Studies Hybridoma Bank (DSHB), and Bloomington Stock Center for antibodies, vectors and fly stocks. Thanks also to the Liverpool Computational Biology Facility and Chris Seidel (Stowers Institute) for assistance with ChIP data analysis, and to the Liverpool Centre for Cell Imaging (https://cci.liv.ac.uk/) for help with microscopy and image analysis. The work was funded by the MRC (MR/K015931/1), NWCR (CR847), Liverpool CRUK Centre and the University of Liverpool international PhD fees waiver scheme.

## References

1. Abdelilah-Seyfried, S., D.N. Cox, and Y.N. Jan. 2003. Bazooka is a permissive factor for the invasive behavior of discs large tumor cells in Drosophila ovarian follicular epithelia. Development. w130:1927–1935.

2. Aguilar-Aragon, M., G. Fletcher, and B.J. Thompson. 2020. The cytoskeletal motor proteins Dynein and MyoV direct apical transport of Crumbs. Dev Biol. 459:126–137.

3. Allan, C., J.M. Burel, J. Moore, C. Blackburn, M. Linkert, S. Loynton, D. Macdonald, W.J. Moore, C. Neves, A. Patterson, M. Porter, A. Tarkowska, B. Loranger, J. Avondo, I. Lagerstedt, L. Lianas, S. Leo, K. Hands, R.T. Hay, A. Patwardhan, C. Best, G.J. Kleywegt, G. Zanetti, and J.R. Swedlow. 2012. OMERO: flexible, model-driven data management for experimental biology. Nat Methods. 9:245–253.

4. Bai, J., and D. Montell. 2002. Eyes absent, a key repressor of polar cell fate during Drosophila oogenesis. Development. 129:5377–5388.

5. Bai, J., Y. Uehara, and D.J. Montell. 2000. Regulation of invasive cell behavior by taiman, a Drosophila protein related to AIB1, a steroid receptor coactivator amplified in breast cancer. Cell. 103:1047–1058.

6. Beccari, S., L. Teixeira, and P. Rorth. 2002. The JAK/STAT pathway is required for border cell migration during Drosophila oogenesis. Mech Dev. 111:115–123.

7. Bianco, A., M. Poukkula, A. Cliffe, J. Mathieu, C.M. Luque, T.A. Fulga, and P. Rorth. 2007. Two distinct modes of guidance signalling during collective migration of border cells. Nature. 448:362–365.

8. Bonnet, J., C.Y. Wang, T. Baptista, S.D. Vincent, W.C. Hsiao, M. Stierle, C.F. Kao, L. Tora, and D. Devys. 2014. The SAGA coactivator complex acts on the whole transcribed genome and is required for RNA polymerase II transcription. Genes Dev. 28:1999–2012.

9. Buracco, S., S. Claydon, and R. Insall. 2019. Control of actin dynamics during cell motility. F1000Res. 8.

10. Burel, J.M., S. Besson, C. Blackburn, M. Carroll, R.K. Ferguson, H. Flynn, K. Gillen, R. Leigh, S. Li, D. Lindner, M. Linkert, W.J. Moore, B. Ramalingam, E. Rozbicki, A. Tarkowska, P. Walczysko, C. Allan, J. Moore, and J.R. Swedlow. 2015. Publishing and sharing multi-dimensional image data with OMERO. Mamm Genome. 26:441–447.

11. Cai, D., S.C. Chen, M. Prasad, L. He, X. Wang, V. Choesmel-Cadamuro, J.K. Sawyer, G. Danuser, and D.J. Montell. 2014. Mechanical feedback through E-cadherin promotes direction sensing during collective cell migration. Cell. 157:1146–1159.

12. Cai, J., M.K. Culley, Y. Zhao, and J. Zhao. 2018. The role of ubiquitination and deubiquitination in the regulation of cell junctions. Protein Cell. 9:754–769.

13. Chen, J.V., L.R. Kao, S.C. Jana, E. Sivan-Loukianova, S. Mendonca, O.A. Cabrera, P. Singh, C. Cabernard, D.F. Eberl, M. Bettencourt-Dias, and T.L. Megraw. 2015. Rootletin organizes the ciliary rootlet to achieve neuron sensory function in Drosophila. J Cell Biol. 211:435–453.

14. Clague, M.J., I. Barsukov, J.M. Coulson, H. Liu, D.J. Rigden, and S. Urbe. 2013. Deubiquitylases from genes to organism. Physiol Rev. 93:1289–1315.

15. Cloud, V., A. Thapa, P. Morales-Sosa, T.M. Miller, S.A. Miller, D. Holsapple, P.M. Gerhart, E. Momtahan, J.L. Jack, E. Leiva, S.R. Rapp, L.G. Shelton, R.A. Pierce, S. Martin-Browns, L. Florens, M.P. Washburn, and R.D. Mohan. 2019. Ataxin-7 and Non-stop coordinate SCAR protein levels, subcellular localization, and actin cytoskeleton organization. Elife. 8.

16. Duchek, P., and P. Rorth. 2001. Guidance of cell migration by EGF receptor signaling during Drosophila oogenesis. Science. 291:131–133.

17. Duchek, P., K. Somogyi, G. Jekely, S. Beccari, and P. Rorth. 2001. Guidance of cell migration by the Drosophila PDGF/VEGF receptor. Cell. 107:17–26.

18. Fernandez, B.G., P. Gaspar, C. Bras-Pereira, B. Jezowska, S.R. Rebelo, and F. Janody. 2011. Actin-Capping Protein and the Hippo pathway regulate F-actin and tissue growth in Drosophila. Development. 138:2337–2346.

19. Fletcher, G.C., E.P. Lucas, R. Brain, A. Tournier, and B.J. Thompson. 2012. Positive feedback and mutual antagonism combine to polarize Crumbs in the Drosophila follicle cell epithelium. Curr Biol. 22:1116–1122.

20. Fulga, T.A., and P. Rorth. 2002. Invasive cell migration is initiated by guided growth of long cellular extensions. Nat Cell Biol. 4:715–719.

21. Gel, B., and E. Serra. 2017. karyoploteR: an R/ Bioconductor package to plot customizable genomes displaying arbitrary data. Bioinformatics. 33:3088–3090.

22. Glinsky, G.V., O. Berezovska, and A.B. Glinskii. 2005. Microarray analysis identifies a death-from-cancer signature predicting the rapy failure in patients with multiple types of cancer. J Clin Invest. 115:1503–1521.

23. Godt, D., and U. Tepass. 2009. Breaking a temporal barrier: signalling crosstalk regulates the initiation of border cell migration. Nat Cell Biol. 11:536–538.

24. Grevengoed, E.E., D.T. Fox, J. Gates, and M. Peifer. 2003. Balancing different types of actin polymerization at distinct sites: roles for Abelson kinase and Enabled. J Cell Biol. 163:1267–1279.

25. Haeger, A., K. Wolf, M.M. Zegers, and P. Friedl. 2015. Collective cell migration: guidance principles and hierarchies. Trends Cell Biol. 25:556–566.

26. Helmlinger, D., S. Marguerat, J. Villen, S.P. Gygi, J. Bahler, and F. Winston. 2008. The S. pombe SAGA complex controls the switch from proliferation to sexual differentiation through the opposing roles of its subunits Gcn5 and Spt8. Genes Dev. 22:3184–3195.

27. Helmlinger, D., S. Marguerat, J. Villen, D.L. Swaney, S.P. Gygi, J. Bahler, and F. Winston. 2011. Tra1 has specific regulatory roles, rather than global functions, within the SAGA co-activator complex. EMBO J. 30:2843–2852.

28. Hillmer, R.E., and B.A. Link. 2019. The Roles of Hippo Signaling Transducers Yap and Taz in Chromatin Remodeling. Cells. 8.

29. Jang, A.C., Y.C. Chang, J. Bai, and D. Montell. 2009. Border-cell migration requires integration of spatial and temporal signals by the BTB protein Abrupt. Nat Cell Biol. 11:569–579.

30. Jiang, H., P.H. Patel, A. Kohlmaier, M.O. Grenley, D.G. McEwen, and B.A. Edgar. 2009. Cytokine/Jak/ Stat signaling mediates regeneration and homeostasis in the Drosophila midgut. Cell. 137:1343–1355.

31. Jonchere, V., and D. Bennett. 2013. Validating RNAi phenotypes in Drosophila using a synthetic RNAi-resistant transgene. PLoS One. 8:e70489.

32. Ko, C., Y.G. Kim, T.P. Le, and K.W. Choi. 2016. Twinstar/ cofilin is required for regulation of epithelial integrity and tissue growth in Drosophila. Oncogene. 35:5144–5154.

33. Kosinsky, R.L., F. Wegwitz, N. Hellbach, M. Dobbelstein, A. Mansouri, T. Vogel, Y. Begus-Nahrmann, and S.A. Johnsen. 2015. Usp22 deficiency impairs intestinal epithelial lineage specification in vivo. Oncotarget. 6:37906–37918.

34. Koutelou, E., C.L. Hirsch, and S.Y. Dent. 2010. Multiple faces of the SAGA complex. Curr Opin Cell Biol. 22:374–382.

35. Krause, M., and A. Gautreau. 2014. Steering cell migration: lamellipodium dynamics and the regulation of directional persistence. Nat Rev Mol Cell Biol. 15:577–590.

36. Kusch, T., S. Guelman, S.M. Abmayr, and J.L. Workman. 2003. Two Drosophila Ada2 homologues function in different multiprotein complexes. Mol Cell Biol. 23:3305–3319.

37. Law, A.L., A. Vehlow, M. Kotini, L. Dodgson, D. Soong, E. Theveneau, C. Bodo, E. Taylor, C. Navarro, U. Perera, M. Michael, G.A. Dunn, D. Bennett, R. Mayor, and M. Krause. 2013. Lamellipodin and the Scar/WAVE complex cooperate to promote cell migration in vivo. J Cell Biol. 203:673–689.

38. Lee, K.K., M.E. Sardiu, S.K. Swanson, J.M. Gilmore, M. Torok, P.A. Grant, L. Florens, J.L. Workman, and M.P. Washburn. 2011. Combinatorial depletion analysis to assemble the network architecture of the SAGA and ADA chromatin remodeling complexes. Mol Syst Biol. 7:503.

39. Lee, T., and L. Luo. 2001. Mosaic analysis with a repressible cell marker (MARCM) for Drosophila neural development. Trends Neurosci. 24:251–254.

40. Lee, T.I., H.C. Causton, F.C. Holstege, W.C. Shen, N. Hannett, E.G. Jennings, F. Winston, M.R. Green, and R.A. Young. 2000. Redundant roles for the TFIID and SAGA complexes in global transcription. Nature. 405:701–704.

41. Li, X., C.W. Seidel, L.T. Szerszen, J.J. Lange, J.L. Workman, and S.M. Abmayr. 2017. Enzymatic modules of the SAGA chromatin-modifying complex play distinct roles in Drosophila gene expression and development. Genes Dev. 31:1588–1600.

42. Lin, T.H., T.H. Yeh, T.W. Wang, and J.Y. Yu. 2014. The Hippo pathway controls border cell migration through distinct mechanisms in outer border cells and polar cells of the Drosophila ovary. Genetics. 198:1087–1099.

43. Lin, Z., H. Yang, Q. Kong, J. Li, S.M. Lee, B. Gao, H. Dong, J. Wei, J. Song, D.D. Zhang, and D. Fang. 2012. USP22 antagonizes p53 transcriptional activation by deubiquitinating Sirt1 to suppress cell apoptosis and is required for mouse embryonic development. Mol Cell. 46:484–494.

44. Lucas, E.P., I. Khanal, P. Gaspar, G.C. Fletcher, C. Polesello, N. Tapon, and B.J. Thompson. 2013. The Hippo pathway polarizes the actin cytoskeleton during collective migration of Drosophila border cells. J Cell Biol. 201:875–885.

45. Margolis, J., and A. Spradling. 1995. Identification and behavior of epithelial stem cells in the Drosophila ovary. Development. 121:3797–3807.

46. Martin, K.A., B. Poeck, H. Roth, A.J. Ebens, L.C. Ballard, and S.L. Zipursky. 1995. Mutations disrupting neuronal connectivity in the Drosophila visual system. Neuron. 14:229–240.

47. McDonald, J.A., A. Khodyakova, G. Aranjuez, C. Dudley, and D.J. Montell. 2008. PAR-1 kinase regulates epithelial detachment and directional protrusion of migrating border cells. Curr Biol. 18:1659–1667.

48. McDonald, J.A., E.M. Pinheiro, and D.J. Montell. 2003. PVF1, a PDGF/VEGF homolog, is sufficient to guide border cells and interacts genetically with Taiman. Development. 130:3469–3478.

49. Medina, E., J. Williams, E. Klipfell, D. Zarnescu, G. Thomas, and A. Le Bivic. 2002. Crumbs interacts with moesin and beta(Heavy)-spectrin in the apical membrane skeleton of Drosophila. J Cell Biol. 158:941–951.

50. Mishra, A.K., J.P. Campanale, J.A. Mondo, and D.J. Montell. 2019. Cell interactions in collective cell migration. Development. 146.

51. Montell, D.J., P. Rorth, and A.C. Spradling. 1992. slow border cells, a locus required for a developmentally regulated cell migration during oogenesis, encodes Drosophila C/EBP. Cell. 71:51–62.

52. Montell, D.J., W.H. Yoon, and M. Starz-Gaiano. 2012. Group choreography: mechanisms orchestrating the collective movement of border cells. Nat Rev Mol Cell Biol. 13:631–645.

53. Muratoglu, S., S. Georgieva, G. Papai, E. Scheer, I. Enunlu, O. Komonyi, I. Cserpan, L. Lebedeva, E. Nabirochkina, A. Udvardy, L. Tora, and I. Boros. 2003. Two different Drosophila ADA2 homologues are present in distinct GCN5 histone acetyltransferase-containing complexes. Mol Cell Biol. 23:306–321.

54. Niewiadomska, P., D. Godt, and U. Tepass. 1999. DE-Cadherin is required for intercellular motility during Drosophila oogenesis. J Cell Biol. 144:533–547.

55. Norden, C., and V. Lecaudey. 2019. Collective cell migration: general themes and new paradigms. Curr Opin Genet Dev. 57:54–60.

56. Pahi, Z., Z. Kiss, O. Komonyi, B.N. Borsos, L. Tora, I.M. Boros, and T. Pankotai. 2015. dTAF10- and dTAF10b-Containing Complexes Are Required for Ecdysone-Driven Larval-Pupal Morphogenesis in Drosophila melanogaster. PLoS One. 10:e0142226.

57. Pankotai, T., O. Komonyi, L. Bodai, Z. Ujfaludi, S. Muratoglu, A. Ciurciu, L. Tora, J. Szabad, and I. Boros. 2005. The homologous Drosophila transcriptional adaptors ADA2a and ADA2b are both required for normal development but have different functions. Mol Cell Biol. 25:8215–8227.

58. Pankotai, T., N. Zsindely, E.E. Vamos, O. Komonyi, L. Bodai, and I.M. Boros. 2013. Functional characterization and gene expression profiling of Drosophila melanogaster short dADA2b isoform-containing dSAGA complexes. BMC Genomics. 14:44.

59. Pinheiro, E.M., and D.J. Montell. 2004. Requirement for Par-6 and Bazooka in Drosophila border cell migration. Development. 131:5243–5251.

60. Plutoni, C., S. Keil, C. Zeledon, L.E.A. Delsin, B. Decelle, P.P. Roux, S. Carreno, and G. Emery. 2019. Misshapen coordinates protrusion restriction and actomyosin contractility during collective cell migration. Nat Commun. 10:3940.

61. Poeck, B., S. Fischer, D. Gunning, S.L. Zipursky, and I. Salecker. 2001. Glial cells mediate target layer selection of retinal axons in the developing visual system of Drosophila. Neuron. 29:99–113.

62. Poukkula, M., A. Cliffe, R. Changede, and P. Rorth. 2011. Cell behaviors regulated by guidance cues in collective migration of border cells. J Cell Biol. 192:513–524.

63. Prasad, M., A.C. Jang, M. Starz-Gaiano, M. Melani, and D.J. Montell. 2007. A protocol for culturing Drosophila melanogaster stage 9 egg chambers for live imaging. Nat Protoc. 2:2467–2473.

64. Quinlan, A.R., and I.M. Hall. 2010. BEDTools: a flexible suite of utilities for comparing genomic features. Bioinformatics. 26:841–842.

65. Ramel, D., X. Wang, C. Laflamme, D.J. Montell, and G. Emery. 2013. Rab11 regulates cell-cell communication during collective cell movements. Nat Cell Biol. 15:317–324.

66. Sansores-Garcia, L., W. Bossuyt, K. Wada, S. Yonemura, C. Tao, H. Sasaki, and G. Halder. 2011. Modulating F-actin organization induces organ growth by affecting the Hippo pathway. EMBO J. 30:2325–2335.

67. Schumacher, L. 2019. Collective Cell Migration in Development. Adv Exp Med Biol. 1146:105–116.

68. Sherrard, K.M., and R.G. Fehon. 2015. The transmembrane protein Crumbs displays complex dynamics during follicular morphogenesis and is regulated competitively by Moesin and aPKC. Development. 142:1869–1878.

69. Silver, D.L., and D.J. Montell. 2001. Paracrine signaling through the JAK/STAT pathway activates invasive behavior of ovarian epithelial cells in Drosophila. Cell. 107:831–841.

70. Stuelten, C.H., C.A. Parent, and D.J. Montell. 2018. Cell motility in cancer invasion and metastasis: insights from simple model organisms. Nat Rev Cancer. 18:296–312.

71. Swatek, K.N., and D. Komander. 2016. Ubiquitin modifications. Cell Res. 26:399–422.

72. Wang, H., Z. Qiu, Z. Xu, S.J. Chen, J. Luo, X. Wang, and J. Chen. 2018. aPKC is a key polarity determinant in coordinating the function of three distinct cell polarities during collective migration. Development. 145.

73. Weake, V.M., J.O. Dyer, C. Seidel, A. Box, S.K. Swanson, A. Peak, L. Florens, M.P. Washburn, S.M. Abmayr, and J.L. Workman. 2011. Posttranscription initiation function of the ubiquitous SAGA complex in tissue-specific gene activation. Genes Dev. 25:1499–1509.

74. Weake, V.M., K.K. Lee, S. Guelman, C.H. Lin, C. Seidel, S.M. Abmayr, and J.L. Workman. 2008. SAGA-mediated H2B deubiquitination controls the development of neuronal connectivity in the Drosophila visual system. EMBO J. 27:394–405.

75. Weake, V.M., and J.L. Workman. 2008. Histone ubiquitination: triggering gene activity. Mol Cell. 29:653–663.

76. Xu, T., and G.M. Rubin. 1993. Analysis of genetic mosaics in developing and adult Drosophila tissues. Development. 117:1223–1237.

77. Zallen, J.A., Y. Cohen, A.M. Hudson, L. Cooley, E. Wieschaus, and E.D. Schejter. 2002. SCAR is a primary regulator of Arp2/3-dependent morphological events in Drosophila. J Cell Biol. 156:689–701.

78. Zhang, X.Y., M. Varthi, S.M. Sykes, C. Phillips, C. Warzecha, W. Zhu, A. Wyce, A.W. Thorne, S.L. Berger, and S.B. McMahon. 2008. The putative cancer stem cell marker USP22 is a subunit of the human SAGA complex required for activated transcription and cell-cycle progression. Mol Cell. 29:102–111.

79. Zsindely, N., T. Pankotai, Z. Ujfaludi, D. Lakatos, O. Komonyi, L. Bodai, L. Tora, and I.M. Boros. 2009. The loss of histone H3 lysine 9 acetylation due to dSAGA-specific dAda2b mutation influences the expression of only a small subset of genes. Nucleic Acids Res. 37:6665–6680.

